# Hierarchical modelling of functional brain networks in population and individuals from big fMRI data

**DOI:** 10.1101/2021.02.01.428496

**Authors:** Seyedeh-Rezvan Farahibozorg, Janine D Bijsterbosch, Weikang Gong, Saad Jbabdi, Stephen M Smith, Samuel J Harrison, Mark W Woolrich

**Author notes:** Last three authors contributed equally (Stephen M Smith, Samuel J Harrison, Mark W Woolrich).

## Abstract

A major goal of large-scale brain imaging datasets is to provide resources for investigating heterogeneous populations. Characterisation of functional brain networks for individual subjects from these datasets will have an enormous potential for prediction of cognitive or clinical traits. We propose for the first time a technique, Stochastic Probabilistic Functional Modes (sPROFUMO), that is scalable to UK Biobank (UKB) with expected 100,000 participants, and hierarchically estimates functional brain networks in individuals and the population, while allowing for bidirectional flow of information between the two. Using simulations, we show the model’s utility, especially in scenarios that involve significant cross-subject variability, or require delineation of fine-grained differences between the networks. Subsequently, by applying the model to resting-state fMRI from 4999 UKB subjects, we mapped resting state networks (RSNs) in single subjects with greater detail than has been possible previously in UKB (>100 RSNs), and demonstrate that these RSNs can predict a range of sensorimotor and higher-level cognitive functions. Furthermore, we demonstrate several advantages of the model over independent component analysis combined with dual-regression (ICA-DR), particularly with respect to the estimation of the spatial configuration of the RSNs and the predictive power for cognitive traits. The proposed model and results can open a new door for future investigations into individualised profiles of brain function from big data.

**Highlights:**

- We introduce stochastic PROFUMO (sPROFUMO) for inferring functional brain networks from big data
- sPROFUMO hierarchically estimates fMRI networks for the population and every individual
- We characterised high dimensional resting state fMRI networks from UK Biobank
- Model outperforms ICA and dual regression for estimation of individual-specific network topography
- We demonstrate the model’s utility for predicting cognitive traits, and capturing subject variability in network topographies versus connectivity

## 1 Introduction

Spontaneous fluctuations in human brain activity, and their interpretations, have been a key focus of human neuroscience research for several years (Biswal et al., 1995; Buckner and Vincent, 2007; Calhoun et al., 2008; Damoiseaux et al., 2006; Raichle et al., 2001; Smith et al., 2013b). Resting state networks (RSNs) characterise functionally synchronised regions that underlie the brain function in the absence of active tasks, and have been replicated in different population cohorts (Fransson et al., 2007; Lee et al., 2013) and using several imaging modalities such as functional Magnetic Resonance Imaging (fMRI) (Allen et al., 2014; Beckmann et al., 2005; Power et al., 2014) and Magneto-/Electroencephalography (Brookes et al., 2011; de Pasquale et al., 2010; Mantini et al., 2007; Vidaurre et al., 2018b), and have been shown to coactivate with spontaneous replay of recently acquired information (Higgins et al., 2021). Large-scale neuroimaging datasets such as the Human Connectome Project (HCP) (Smith et al., 2013a; Van Essen et al., 2012) and UK Biobank (UKB) (Alfaro-Almagro et al., 2018; Miller et al., 2016), have significantly advanced RSN research, leading to the mapping of brain function with unprecedented detail (Fan et al., 2016; Glasser et al., 2016). Furthermore, the wealth of non-imaging phenotypes in these datasets has provided new insights into the translational importance of the RSNs, which display significant associations with life factors, genetics, behavioural and clinical traits (Elliott et al., 2018; Finn et al., 2015; Jiang et al., 2020; Kong et al., 2019; Vidaurre et al., 2017). What remains largely unresolved, however, is how to accurately and robustly model cross-individual variations of the RSNs in big epidemiological data, such that substantial degrees of population heterogeneity are interpretably accounted for (Smith et al., 2013b). This is particularly important if we are interested in utilising RSNs to characterise cognitive idiosyncrasies in individuals or as biomarkers to predict, e.g., pathology before clinical onset.

Functional brain modes^1^in rest and task are conventionally modelled using group-average algorithms, such as independent component analysis (ICA) (McKeown et al., 1998). Typically, modes are modelled as spatially contiguous parcels (Bellec et al., 2010; Craddock et al., 2012; van den Heuvel et al., 2008) or functionally unified systems distributed over multiple brain areas (Beckmann and Smith, 2004; Calhoun and Adali, 2012; Thomas Yeo et al., 2011), and characterised in terms of spatial configuration over the brain voxels (mode topography) and a summary time course that captures mode activity over time. Of particular interest is to characterise the functional connectivity between the modes themselves, ideally using models that can accurately disambiguate changes in functional connectivity across separate dimensions (e.g. spatial or temporal features) (Bijsterbosch et al., 2019, 2018), thereby obtaining functional connectomes.

Recent evidence indicates that even after registration of the subject data to a standard brain such as MNI space, functional modes still significantly vary across individuals (Glasser et al., 2016; Gordon et al., 2017a; Mueller et al., 2013). These misalignments can be due to multiple factors including limitations in methodologies for aligning subjects’ data (e.g. registration errors), differences in subjects’ brain anatomy (Devlin and Poldrack, 2007; Llera et al., 2019) or inherent differences in functional localisations of the modes (Bijsterbosch et al., 2018; Gordon et al., 2017b; Haxby et al., 2020; Laumann et al., 2015). Therefore, it is increasingly desirable to devise models with hierarchical links between the population and individuals, which can accurately cope with subject deviations from the group, while at the same time maintaining a representative group model that provides correspondence over subjects. Hierarchical models can differ with respect to multiple factors contributing to subject and/or group level mode estimations. Some of the key factors include: the direction of information flow between population and individuals; explicit versus derivational mode estimations for individuals; iterative versus one-off mode estimations; defining hierarchies over mode topography or functional connectivity; defining modes as distributed networks versus contiguous parcels; hard versus soft boundaries between modes; and scaling of algorithms to modern big data such as HCP (∼1000 subjects) and UKB (∼100,000 subjects). Details of these factors and some examples in existing models are presented in Appendix A.

A key element distinguishing hierarchical models is the way the direction of information flow between the population and individuals is set up. Unidirectional models often start from group-level estimations and derive subject modes as a variant of the group, of which dual regression is a well-known example (Beckmann et al., 2009; Nickerson et al., 2017). It runs a two-step multiple regression paradigm: firstly, group maps, typically from spatial ICA, are regressed against the data to extract subject-specific fMRI time series; secondly, DR regresses subject mode time courses against fMRI time-series to obtain subject-specific spatial mode maps. Alternatively, bidirectional models such as multi-subject dictionary learning (Abraham et al., 2013; Varoquaux et al., 2011), hierarchical ICA (Guo and Tang, 2013; Shi and Guo, 2016), hierarchical topographic factor analysis (Manning et al., 2018) and the framework of Probabilistic Functional Modes (PROFUMO) (Harrison et al., 2015) allow for bottom-up, data-driven estimation of the group-level modes, as well as top-down regularisation of the subject specific modes using the group-level model - and can thus accommodate larger degrees of population heterogeneity.

Another important factor that distinguishes different hierarchical models is whether the hierarchy is defined over the mode topography or functional connectivity. As elaborated in Appendix A, existing algorithms often define hierarchy on spatial maps, thus identifying a consensus spatial layout for the modes, and estimating how the spatial arrangement of the voxels belonging to the mode varies across individuals (Glasser et al., 2016; Manning et al., 2018; Mejia et al., 2019; Nickerson et al., 2017; Shi and Guo, 2016). An alternative approach is to estimate a between-mode functional connectivity matrix (i.e. NetMats) at the group-level, and hierarchically estimate subject-specific connectivity through making comparisons with the group (Chong et al., 2017). Notably, however, defining hierarchy solely on spatial topography or temporal connectivity might neglect how these two elements interact with each other in reality, which might in turn result in topographical misalignments being (mis)interpreted as (being changes in) functional connectivity or vice versa (Bijsterbosch et al., 2018). One solution is proposed by the latest version of PROFUMO (Harrison et al., 2020) which defines hierarchical models on both spatial topographies and functional connectivity.

Despite their promise for modelling population heterogeneity, the application of hierarchical models to modern high-resolution fMRI with thousands of subjects is limited because of computational costs. Recent work has proposed solutions that are applicable to group models or unidirectional hierarchical models, e.g. using incremental or stochastic matrix factorisations (Mensch et al., 2018; Smith et al., 2014), or utilising Bayesian priors from group estimations to inform subject-specific network modelling (Mejia et al., 2019). Nevertheless, the application of the bidirectional hierarchical models to big data remains impractical (c.f. Figure 1) (Mejia et al., 2019).

**Figure 1.**
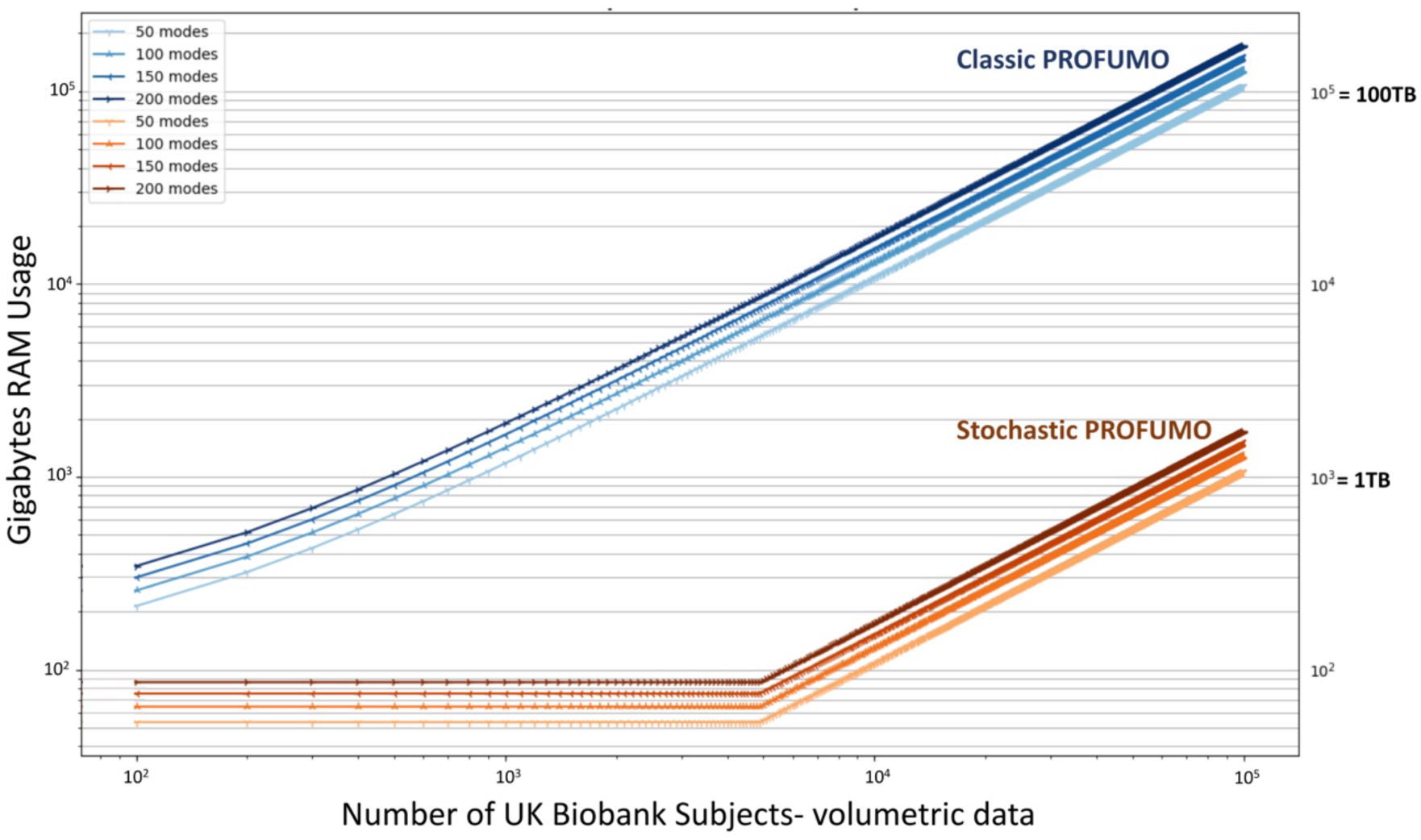
Computational efficiency of Stochastic PROFUMO: CPU RAM required to run the model on up to 100,000 participants, compared to classic PROFUMO. RAM usage for stochastic PROFUMO is calculated with batch size set to 50 or 1% of the full population, whichever the larger. Here we have used volumetric data from UK Biobank as reference for the calculations, which includes ∼230,000 voxels (brain masked), 1 recording session and 490 timepoints per recording. Both x- and y-axes are logarithmic scales.

Contributions of the current study are twofold. Firstly, we propose an advance to PROFUMO using *stochastic* variational Bayes (VB) that can reduce the computational costs by a factor of 100, in order to scale the model for big data. We refer to the proposed model as stochastic PROFUMO, or sPROFUMO for short, and refer to the probabilistic functional modes identified using this approach as PFMs. Stochastic VB (Hoffman et al., 2013) entails partitioning small, random subsets of observations into numerous batches, and visiting only a fraction of the population in every iteration, thus significantly reducing the computational expense (Figure 1), without loss of generalisability. For the purposes of fMRI analyses, different subjects provide a natural way to subdivide the data, and our algorithm therefore iterates through updating the group model over batches, while simultaneously optimising subject modes within these batches. This approach has been successfully applied to neuroimaging data to discover functional modes using group-level Hidden Markov Models (HMM) (Vidaurre et al., 2018a). Generalising this approach in sPROFUMO requires modifications to the original algorithm to ensure preserving group-level generalisability, cross-subject variability and subject-group alignment, as we will elaborate later in section Model. In addition to scaling the model to work on big data, sPROFUMO substantially reduces the computational expenses for medium-sized datasets while preserving the accuracy of estimations, and thus is expected to facilitate the model’s broader usability.

The second contribution of this study is to unravel the cognitive relevance of sPROFUMO RSNs in individual subjects, and disentangle the spatial versus temporal properties of RSNs that contribute to the prediction of cognitive function. We applied sPROFUMO to fMRI data from 4999 UKB subjects, which allowed us to obtain both our first high-dimensional PFM decomposition of brain function (100, 150 and 200 modes), and an increase in the detail of mapping of RSNs in single subjects compared to what has been reported previously from UKB. We tested how accurately these PFMs can predict 68 cognitive scores that cover a range of sensorimotor, memory, executive functions and general fluid intelligence. Our focus on high-dimensional RSNs was informed by recent studies that suggest the functional parcellations with 100-1000 modes to provide more elaborate delineation of the brain function compared to low-dimensional decompositions, which is useful for prediction of non-imaging variables (Dadi et al., 2020; Pervaiz et al., 2020). We investigated in detail the prediction power of mode elements in spatial and temporal domains, and identified the sPROFUMO modes, or PFMs, that provided the best predictors. We further carried out a detailed comparison between sPROFUMO and ICA-DR with regards to accounting for cross-subject variability and prediction power for cognitive heterogeneity.

## 2 Model

In this section, we first provide a brief conceptual summary of PROFUMO (see (Harrison et al., 2020) for further details), and next elaborate on the stochastic variational inference and its application to obtain sPROFUMO.

### 2.1 Classic PROFUMO summary

PROFUMO is a hierarchical matrix factorisation framework with two levels of subject and group modelling. At the subject level, fMRI timeseries (**D**^sr^) are decomposed into a set of spatial maps (**P**^s^), time courses (**A**^sr^) and time course amplitudes (**H**^sr^), with residuals **ε**^sr^:

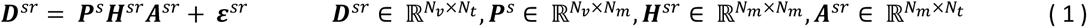

where s denotes each subject and r denotes a recording session. **P**^s^denotes spatial mode layout of the modes across the brain voxels, which hereafter we will refer to as *spatial maps*, or just *maps*. **A**^sr^represents mode time courses and**H**^sr^denotes mode amplitudes. N_v_,N_m_ andN_t_ denote the number of voxels, modes and time points, respectively.

Assuming that anatomical and functional organisation of the brain regions remain unchanged for a subject, the spatial map, **P**^s^, are assumed to be the same across multiple scans (or runs – sets of continuous timeseries data). Each map is modelled using a Double-Gaussian Mixture Model (DGMM), where one Gaussian component is set to account for the signal and a second Gaussian distribution captures the background spatial noise in each voxel.

Additionally, a per-mode membership probability is estimated to determine the probability of a voxel belonging to the signal versus the noise component. Therefore, **P**^s^consists of five posterior parameters (one map for each): signal mean, signal variance, noise mean, noise variance and membership probability. Spatial maps are one of the elements that are estimated hierarchically in PROFUMO. Therefore, in addition to **P**^s^for each subject, a consensus set of group-level parameters is also estimated to capture both the group maps and the key patterns of cross-subject variability in the spatial domain. Concretely, we introduce a set of hyperpriors on - and infer the group-level posteriors over - the signal means, signal variances, noise variances and membership probabilities.

The mode time courses, **A**^sr^, are estimated separately for each scan. These timeseries are allowed to vary across runs such that PROFUMO can account for both the unconstrained nature of the resting state and the inherent cross-run variations in task data. Similar to the spatial maps, mode time courses are also modelled as a sum of a signal and a noise component, where the signal component is HRF-constrained and the noise component follows a Gaussian distribution. **A**^sr^ is not modelled hierarchically, because in general (at least for resting state data) there will be no consensus temporal structure across the subjects. However, even though the time courses are unconstrained, the temporal correlations among the neural time courses can be both highly structured within-individuals, and have a consensus structure across individuals (Shehzad et al., 2009). In order to account for these correlations, PROFUMO models the precision matrix **α**^sr^hierarchically and using Wishart distributions. Additionally, because of haemodynamic responses that govern the BOLD signal, the estimation of the partial correlation matrices explicitly incorporates HRF-constrained autocorrelations between the modes.

Finally, the mode amplitudes, **H**^sr^, are modelled as positive diagonal matrices, to capture (subject and/or scan) variations in the amplitude of the mode time series. In the dual regression paradigm, these amplitudes appear as the time course standard deviations and have been shown to reflect meaningful sources of within- and between-subject variations (Bijsterbosch et al., 2017). The PROFUMO framework captures these variations via multivariate normal distributions, again with a hierarchical link between the group and the subject levels.

### 2.2 Stochastic PROFUMO (sPROFUMO)

#### 2.2.1 Standard VB

PROFUMO uses a variational Bayesian (VB) framework to find a solution for the full probabilistic model described in the previous section. This optimises the parameters of a simple approximating distribution **q**, with the aim that it is as close as possible to the true posterior. In each model iteration, the group model is used to constrain the subject-specific matrix factorisations, from which one can infer posterior distributions for subject-specific spatial maps, time course correlations and amplitudes. Next, the posterior evidence is accumulated and combined across individuals to obtain an updated version of the group model. The model iterates between these two levels of estimation until convergence. In order to reach convergence, the algorithm utilises the conjugate-exponential structure of the model to obtain a closed-form version of the natural gradient of the Free Energy (i.e. the difference between the true and approximate posteriors, as measured by the Kullback-Leibler (KL) divergence), based on which we optimise via gradient ascent. Therefore, for observations ***D*** (where ***D*** is a collection of all ***D****^sr^*in Equation 1), group-level latent variables **λ** and subject-level latent variables ***z***, we will have:

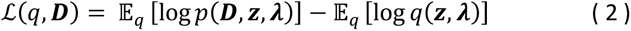

where 𝔼_q_ [log *p*(***D***, ***z***, **λ**)] is the expected log joint, log **q**(***z***, **λ**) is the (approximate) log joint posterior distribution over the latent variables, and ℒ(**q**, ***D***)is the free energy of the model. To find a (locally) optimal solution, traditional VB optimisation must have access to the entire dataset in each iteration, which becomes computationally intractable when a large number of subjects need to be analysed.

#### 2.2.2 Substitution of standard VB with stochastic VB to obtain sPROFUMO

One of the main contributions of this work is to reduce the computational expense of the VB optimisation, by substituting standard VB with stochastic VB (Hoffman et al., 2013). Applying this approach to PROFUMO requires modifications to the original algorithm to ensure that: a) the group model is not biased by a few batches and instead remains representative of the entire population, b) single subjects are kept consistent with the group regardless of when they have been visited, and c) the model has flexibility to accommodate single-subject heterogeneity in a big data population.

We start by making use of the fact that in VB the group- and subject-level PROFUMO parameters are inferred independently, and define them as global and local variables, respectively. Specifically, mean-field approximations allow for factorisation of the joint posteriors into a global term and a product of local terms; i.e.:

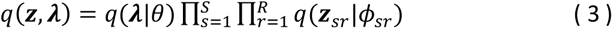

where **λ**, ***z*** are global (i.e. group) and local (i.e. subject) latent variables respectively, governed by global and local parameters *θ* and *ϕ*. Here, S and R denote the total number of subjects and recordings per subject. The mean-field approximations allow for simplification of the free energy estimations into a sum of a global and a local term such that:

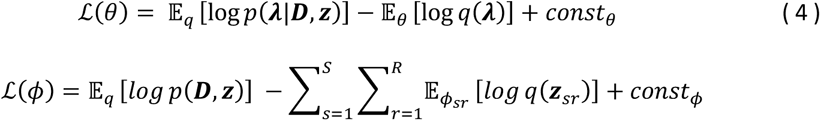

Classic VB iterates between keeping *θ* fixed and optimising with respect to *ϕ =>* ∇ *_ϕ_* ℒ(*ϕ*)= 0 and keeping *ϕ* fixed and optimising with respect to *θ =>* ∇ *_ϕ_* ℒ(*θ*)= 0 until convergence (where ∇*_ ϕ_* ℒ(*ϕ*) and ∇ _*θ*_ ℒ(*θ*) denote gradients of ℒ with respect to *ϕ* and *θ*, respectively).

With stochastic VB, we instead randomly subsample the local variables (i.e. sessions/subjects) across batches, and focus on continuously improving the estimation of the global parameters as more observations are visited over time. Considering ℒ_I_ (*θ*)as the free energy term corresponding to the I^th^ batch, we will have:

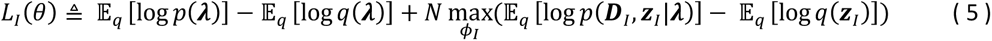

The third term on the right-hand side refers to the optimisation of the local variables within batch I, while the overall aim will be to optimise *L*_I_(*θ*) across batches. The natural gradient of *L*_I_ (*θ*) is therefore a noisy estimate of the natural gradient of the overall variational objective. According to proofs provided by Hoffman *et al*. (2013), even though the noisy gradient 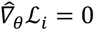 does not have an analytical solution, the conjugate-exponential structure of PROFUMO means that convergence can be achieved if: a) we scale the subject-specific terms in Equation 5 in each batch by the factor N (i.e. number of batches), such that *L*_I_(*θ*) is updated as though the entire population was included; and b) global parameters are updated iteratively, using a weighted sum of the intermediate global parameters obtained from the current batch 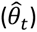 and what has been accumulated over all the previous batches (*θ_t_*_-1_). In other words, we keep a memory of the global parameter updates over time; i.e.:

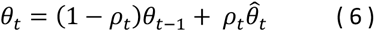

We adopt a similar approach as (Hoffman et al., 2013; Vidaurre et al., 2018a), in defining a decreasing step size for the stochastic gradient descent:

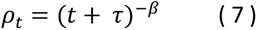

τ > 0 is the delay parameter of stochastic inference that determines the degree to which initial batches influence the overall inference, and β ∈ (0.5, 1] is the forget rate of the model that determines the degree to which global parameter updates rely on the current batch versus previous batches (i.e. inter-batch variations). Larger β thus corresponds to smaller ρ_t_ and results in less stochasticity in the model.

Having described the general rationale behind stochastic VB inference and application to PROFUMO, we next elaborate specific changes in the model implementation in sections 2.2.3 to 2.2.5. A flowchart of these steps is outlined in Figure 2.

**Figure 2.**
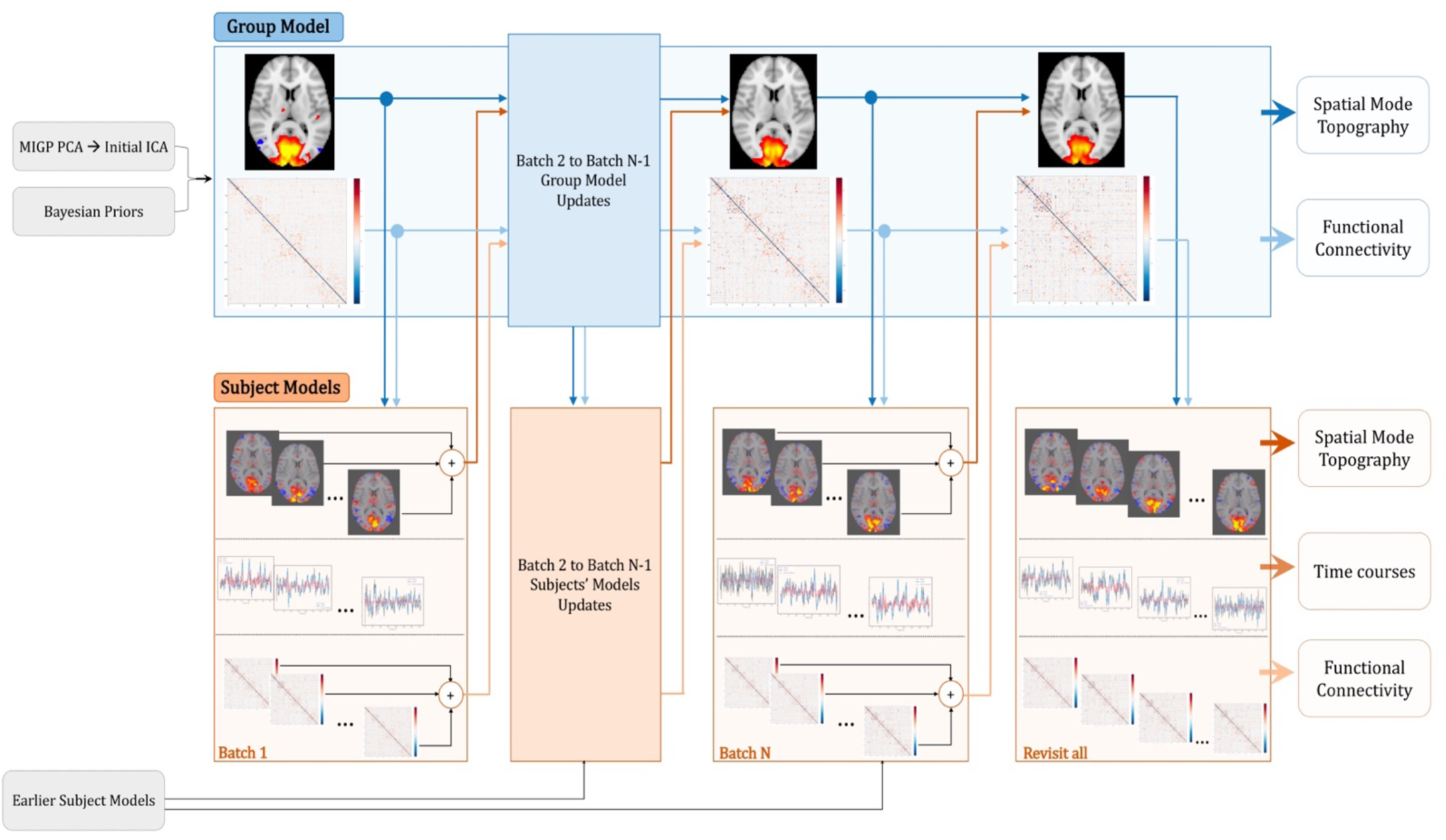
Flowchart of different steps of stochastic PROFUMO (sPROFUMO). Batch updates, progressing from left to right, involve treating group model as global parameters that are continuously updated over time, whilst subsets of subjects are randomly selected and respective parameters are locally optimised within each batch. Blue boxes: group model update process, blue arrows: flow of information from the group to the subjects, where dark blue denotes hierarchical links between the spatial maps and light blue denotes hierarchical links between partial temporal correlation matrices. Orange boxes: subject models and batch-based update process, orange arrows: flow of information from subjects/batches to the group, where dark orange denotes hierarchical links between the spatial maps and light orange denotes hierarchical link between partial temporal correlation matrices. Grey: initialisation and priors, MIGP: MELODIC’s Incremental Group-PCA.

#### 2.2.3 Changes in model initialisation

As outlined in 2.1, PROFUMO estimates multiple parameters per voxel and mode. For a typical volumetric fMRI in 2mm MNI space, this entails estimating of the order of 150,000,000 parameters per subject. Considering that with a model of this complexity, random initialisations are likely not viable, the algorithm initialises the group maps based on a group-level variational ICA in order to place the parameters within a realistic ballpark. To obtain this initialisation, subject fMRI data matrices **D**^sr^are first dimension reduced using SVD to obtain **X**^sr^∈ ℝ*^N^_v_^xN^_p_* where p is the number of top singular values. The **X**^sr^ matrices are concatenated across subjects and further reduced using random SVD to 2*N_m_ spatial basis vectors.

However, to handle many thousands of subjects, e.g. in UKB, the SVD approach used in the original PROFUMO will become intractable as it requires concatenation of the subject data from the entire population in one matrix. We therefore substituted this step in sPROFUMO with MELODIC’s Incremental Group-PCA (MIGP) (Smith et al., 2014). MIGP is an online PCA decomposition, where subjects’ data are visited one at a time, and dimension reduced to 2*N_m_ in every step, until the entire population is visited. Considering that MIGP loads only a few subjects’ data at a time, it offers an additional advantage of allowing us to work with the full rank subject data **D**^sr^ instead of dimension-reduced **X**^sr^, and is thus expected to preserve fine-grained modes that are likely to get eliminated in dimension reductions.

After obtaining the spatial basis, group-level variational spatial ICA (Harrison et al., 2020; Lawrence and Bishop, 2000) is applied to the PCA output in order to infer N_m_ initial maps. These initial maps, together with hyperpriors, are used to initialise the group model in PROFUMO/sPROFUMO. Note that initial maps do not provide subject-specific results, subject-specific decompositions are conducted during full inference, with the group model used as a prior.

#### 2.2.4 Batch randomisations and changes in update rules

Considering that we randomise subjects across numerous small batches of size K, and in order to maximise the chances of every subject being selected multiple times, the randomisation is weighted by a parameter 𝑤*_s_*_”_ = τ*^ni^* that determines the probability of a subject i being picked in the current batch, where n_i_ denotes the number of times that the subject i has been picked so far. Subjects that are picked for a batch are initialised as follows:

- When a subject is visited for the first time, we can either initialise them based on the initial maps (from variational ICA as outlined earlier in 2.2.3) or the latest group model. There are arguably pros and cons to each approach; namely, initialisation based on the latest group will align the subjects and can thus help with quicker model convergence. However, considering that any subsequent batches will be built upon the first few batches, it might result in the overall inference being biased towards the early batches, thus potentially compromising the representativeness of the final group model. We therefore chose to always initialise subjects’ first visit based on the initial maps.
- In any subsequent visits, subject-specific updates build upon previous incarnations of the subject model. In this way, by keeping a memory of a subject’s own previous visits, it can be expected that subject-specific deviations from the group are more accurately accounted for.
- After subject-initialisations, we initially keep the group model constant and run multi-iterations of subject updates to bring the subjects into better alignment with the group. We refer to this step as *initial* batch updates.
- Subsequently, we run the full inference (section 2.1) on subjects within current batch (local parameters in stochastic VB) as well as the group model (global parameters). We refer to this step as *full* batch updates, or batch updates for short.

As outlined earlier, we treat group-level parameters as global parameters that stochastic VB optimises continuously across batches. Considering that all the distributions that are used in the model are from the exponential family (e.g. Gaussian), changes in the group-level update rules follow Equation 6.

It is worth noting that sPROFUMO can be expected to allow for larger degrees of heterogeneity; i.e. subject deviations from the group average. This is due to the fact that in PROFUMO, all subjects are updated in line with the group in every model iteration, which will produce a tendency for the algorithm to converge to local optima where the subject-level models are close to the group average. On the contrary, in sPROFUMO we are updating stochastically by using a random subset of the subjects at each model iteration with each subject being only visited at random intervals; and though the model update procedures are designed to ensure subject-group alignment throughout iterations, this stochastic updating may help sPROFUMO to jump out of local minima (Keskar et al., 2017; Kleinberg et al., 2018).

#### 2.2.5 Additional considerations

Batch randomisations mean that some of the subjects will be selected during the earliest batches and may not be revisited later. We therefore revisit every subject again at the end of the inference and update them with reference to the final group model. This is done to ensure the final estimations of posteriors for all subjects are defined in relation to the same group model.

Additionally, with regards to the choice of the stochastic parameters in EQUATIONS 6 and 7, it is typically of interest to minimise the running times and RAM memory requirements of the model, while maximising the accuracies of estimations and reaching a convergence point. The number of batches, and initial and full batch updates, control the running times while the choice of batch size controls the memory usage. It is therefore of interest to find an optimal trade-off between these factors, which we will later explore using simulations.

## 3 Materials and Methods

### 3.1 Simulations

We performed two sets of simulations. The first set focused on evaluating sPROFUMO’s performance compared to PROFUMO, where the entire population is available at every model iteration. More specifically, we tested how the choice of the key stochastic parameters, namely batch size, stochastic forget rate β and full batch updates, affects the accuracy of final results. The second set of simulations were aimed at evaluating sPROFUMO’s performance in obtaining high dimensional modes; and in this case, comparison is with the spatial ICA-dual regression paradigm. We explored several scenarios that make the latter set of simulations challenging, particularly with respect to accurate reconstruction of high dimensional subject-specific networks. More details of each simulation set are presented in the RESULTS section. For each simulation scenario, we created synthetic data with realistic fMRI settings for 500 subjects with two recording sessions for each subject. Each simulation scenario was repeated twice and results were pooled for reporting.

Overall, we investigated data from 11 scenarios, each repeated twice and each repeat consisting of two datasets (2 x 500) of simulated subjects, providing a thorough evaluation of our model. Each synthetic fMRI data matrix consisted of 10,000 voxels and 300 time points at a TR of 0.72s, and was created using an outer product model, similar to Equation 1. Therefore, spatial and temporal properties of the modes are simulated independently in this pipeline.

At the group level, we defined a set of spatial maps **P_g_** that consisted of modes generated from a number of randomly-selected contiguous blocks of voxels (i.e. parcels), such that some of the modes were confined to one brain region and others were distributed over multiple non-contiguous regions. Weights of signal for each voxel within a mode were drawn from a Gamma distribution. Subject-specific variations of these parcels, and subsequently subject spatial mode maps, **P_s_**, were defined at the vicinity of the group maps by applying spatial warps and adding background Gaussian noise. Time courses were generated separately for each mode, subject and session, but we defined a hierarchical link between the group and subjects’ temporal correlation matrices, following a Wishart distribution, to ensure a consensus representation of the functional connectivity between the two levels. Time courses were first generated as semi-Gaussian neural time course with amplified frequencies < 0.1Hz, and next convolved with a random draw from the FLOBS basis functions (Woolrich et al., 2004) such that final time courses would mimic the BOLD signal. Finally, random noise was added to the outer product of spatial maps and time courses to create space-time data matrices. More details of simulation parameters are available in (Bijsterbosch et al., 2019; Harrison et al., 2020, 2015).

The simulated modes were reconstructed using sPROFUMO, PROFUMO and ICA/ICA-DR, and results were compared against the ground truth. For this purpose, we initialised all three models based on the same set of spatial bases that were obtained using MIGP, to ensure that the observed differences are not due to the initial PCA. Additionally, we have initialised PROFUMO and sPROFUMO based on the same set of initial maps.

### 3.2 Applying sPROFUMO to UK Biobank

We used UK Biobank (UKB) data, accessed under application number 8107, for volumetric resting state fMRI data as well as cognitive tests and imaging confounds (see 3.4.1 and 3.4.2 for more details on the latter). We randomly drew 4999 subjects from the May 2019 release of this dataset. fMRI data consisted of one recording session per subject, with 490 (occasionally 530) time points per session, at a TR of 0.735s, yielding ∼6 minutes of data per subject. Data were pre-processed using the standard UKB pipeline that includes quality control, brain extraction, motion correction, artefact rejection using FSL-FIX, high-pass temporal filtering (sigma = 50.0s, Gaussian-weighted least-squares straight line fitting) and registration to standard MNI-2mm space (Alfaro-Almagro et al., 2018). As a part of sPROFUMO’s internal initialisation (see (Harrison et al., 2020) for details), we further processed the data using voxel-wise removal of timecourse mean and variance normalisation. We next initialised the model as explained in 2.2.3 and ran full model inference thorough batch randomisations as outlined in 2.2.4 and 2.2.5, and characterised 100, 150 and 200 sPROFUMO modes.

#### 3.2.1 Dealing with the missing modes to obtain high dimensional decompositions

In previous PROFUMO papers and based on real resting state fMRI data from up to ∼1000 subjects in HCP Young Adult cohort, we reported that while the model reliably estimates a low dimensional set of modes (i.e. up to 40), it tends to eliminate any more modes beyond that (Harrison et al., 2020). Additionally, some of the modes that are reliably estimated in the group might be returned as empty in a subset of subjects. This is despite the fact that ICA and ICA-DR have reported up to 300 modes from the same data (Smith et al., 2015). The mode elimination in PROFUMO corresponds to the lack of evidence – in a Bayesian sense – for the mode being present, given that subject’s data and the inferred noise level. Together, these can render the posterior probabilities of mode presence as zero. The validity of, and reasons behind, these missing modes had remained puzzling to date. More importantly, it is unclear whether these are specific to the s/PROFUMO framework or might also be reflected in a different way in other models e.g. ICA-DR.

In Appendix B we will present details of five main changes in the data and model that were found to largely resolve sPROFUMO missing modes for the UK Biobank data in the current study; these include: 1) increased group-level signal-to-noise ratio due to the higher number of subjects in UKB; 2) *stochastic* variational inference in sPROFUMO may have allowed for the VB optimisation to jump out of local minima, resulting in an improved minima that seems to allow for a higher degrees of subject variability in the group model; 3) homogeneous spatial smoothness in different parts of the brain (e.g. cortical and subcortical regions); 4) no SVD dimensionality reduction at single subject level, and 5) increased subject-level SNR by applying a small amount of spatial smoothing. Factors 1, 2 and 3 led to a higher number of group-level modes being reliably reconstructed and factors 2, 4 and 5 increased the number of subject-specific modes.

### 3.3 Applying spatial ICA and ICA-Dual Regression to UK Biobank

In addition to sPROFUMO characterisation of functional brain modes from resting state fMRI data, we also used spatial ICA followed by dual regression (DR) to characterise the modes. The ICA and ICA-DR are among the most commonly used methods for group-level and subject-level estimation of the resting state networks, respectively. Additionally, ICA/ICA-DR have been used as standard RSN characterisation methods from large-scale datasets such as HCP and UKB. Thus, they can provide suitable models to compare sPROFUMO results against. We used FSL tool *MELODIC* to identify 150 spatially independent ICA modes at the group-level and subsequently used FSL tool *dual_regression* to map the ICA group-level results onto single subject data. Stage 1 of Dual Regression yielded subject-specific time courses for each mode and stage 2 yielded subject-specific spatial maps. We further computed mode amplitudes as standard deviation of the time courses and estimated functional connectivity between the modes using Tikhonov-regularised partial correlations. It is worth noting that in order to make the final sPROFUMO and ICA/ICA-DR results fully comparable, we used the same PCA initialisation for both models. For this purpose, MIGP results were obtained as explained earlier in 2.2.3 and were fed into both MELODIC and sPROFUMO for initialisation.

### 3.4 Prediction pipeline

We investigated the accuracies of the spatial and temporal properties of sPROFUMO modes (i.e. PFMs) in predicting cognitive outcome from behavioural cognitive tests in UKB. We conducted predictions at two levels: multi-mode predictions where output from all the modes were combined, and uni-mode predictions where each mode was used separately for prediction. For both levels, we conducted predictions based on: a) Mode spatial maps, b) Mode temporal network matrices (temporal NetMats; partial correlation matrix between PFM timecourses), c) Mode spatial NetMats (full correlation matrix between PFM maps), d) Mode amplitudes and e) multi-element; i.e. all 4 elements combined. We further compared PFM prediction accuracies to ICA-DR.

#### 3.4.1 Selecting Cognitive Tests

As the first step, we selected a subset of outcome measures from UK Biobank cognitive tests by pruning 1172 metrics to 68. This initial filtering was based on two criteria, firstly, one of the authors manually screened the outcome measures and only included active measures that were directly reflective of each subject’s performance. For example, among outputs from the “Reaction Time” task, which entails viewing two cards (A and B) and pressing buttons if they are identical, we included the “Mean time to correctly identify matches”, while “Index for card A in round” was excluded. Secondly, we only selected tests that had a non-NaN value in at least 25% of the subjects. The final list of 68 tests belonged to the following categories: *Reaction time, Trail making, Matrix pattern completion, Numeric memory, Prospective memory, Pairs matching, Symbol digit substitution* and *Fluid intelligence*. These tests cover a wide range of cognitive abilities from sensory-motor coordination to memory and executive functions, thus providing a suitable testbed for evaluating the cognitive relevance of the PFMs. The list of 68 measures are shown in Table S 1 and more details are available in UKB website: https://biobank.ctsu.ox.ac.uk/crystal/label.cgi?id=100026.

#### 3.4.2 Confound Removal

The large number of subjects in UKB provides an unparalleled potential for application of machine learning algorithms and predicting non-imaging phenotypes based on image-extracted brain features. However, contamination of the imaging data by interfering factors, often referred to as imaging confounds, is likely to have a significant impact on the interpretability of the results (Alfaro-Almagro et al., 2020; Snoek et al., 2019). One solution to alleviate this problem is to regress out the confounds (a.k.a. deconfounding) before applying machine learning predictions. There is not currently a consensus in the literature as to whether or not deconfounding is required in prediction pipelines. Indeed, if the aim is to obtain the highest prediction accuracies for non-imaging phenotypes, regardless of the factors that drive the accuracies, one may opt to run predictions without deconfounding. However, that might arguably interfere with the interpretability of the results in scenarios where a common confounding factor, e.g. head size or head motion might have contributed to prediction accuracies (Snoek et al., 2019). Here, in order to improve the interpretability of our results, we chose to run predictions after deconfounding.

In a recent study, Alfaro-Almagro *et al*. (2020) provided a comprehensive set of 602 confounds in UKB brain imaging. Here we used a reduced set by: a) selecting conventional confounds including age, age squared, sex, age x sex, site, head size and head motion; b) reducing the remaining confounds by applying singular value decompositions and retrieving the top principal components that explained 85% of the variance. These two steps yielded 82 variables for deconfounding, which were regressed out of both predictor and target variables using linear regression. Importantly, we applied deconfounding within cross-validation folds in order to avoid leakage of information from test to train data, as proposed by (Snoek et al., 2019).

#### 3.4.3 Elastic-Net prediction and cross-validation

The prediction pipeline was implemented in Python 3.6.5 using scikit-learn 0.19.1 (Pedregosa et al., 2011). It was built around ElasticNet regression and nested 5-fold cross validations, where subjects were split into 5 non-overlapping subsets, and in each iteration 80% of the subjects were assigned to train and 20% to the test group. In each cross-validation loop, where pre-prediction feature selections were required (details in the following sections), the top n% of the features were selected based on correlation with the target variable within the training set. Next, quantile transformation (*QuantileTransformer*) was applied to obtain a Gaussian distribution for each predictor and target variable across subjects. The quantiles were estimated using the training set and applied to transform both train and test data. Next, confound regression parameters (or “betas”) were estimated from the training set and applied to de-confound both train and test data. Finally, *ElasticNetCV* was used to predict target variables in the test set. Note that we used nested cross-validations within the training set to optimise the ratio of Lasso to Lasso+Ridge regularisation (*l1ratio* varied between [0.1,0.5,0.7,0.9,0.95,0.99,1.0] with 10 alphas per *l1ratio*). The prediction accuracies were calculated based on correlations between estimated and actual values of the target variables across subjects in the test set.

#### 3.4.4 Multi-mode prediction

Multi-mode prediction involved combining output from all the modes to predict cognitive outcome based on sPROFUMO and ICA-DR modes:

- For the spatial maps, estimating 150 modes resulted in a **P**_grand_ matrix of N_subject_ x N_voxel_ x 150, yielding ∼35 million features per subject. Dimensionality reduction of this matrix to a few hundred features that can meaningfully capture the essence of subject spatial maps is non-trivial. For this purpose, we used unsupervised learning in the form of FMRIB’s Linked ICA for big data (bigFLICA) (Gong et al., 2021; Groves et al., 2011), which is an ICA framework originally proposed to fuse multimodal data. First, we collapsed subject spatial maps across voxels using sparse dictionary learning and obtained a **P**_dicL_ matrix of size N_subject_ x 1000 x 150. Next, we considered each mode as a separate “modality” within bigFLICA and obtained a **P**_feat_ feature matrix of size N_subject_ x 500. Using linked ICA in this way will allow us to preserve subject-specific variations in each mode’s spatial map and combine them across numerous modes to obtain a set of independent features to characterise the sources of population variations in PFMs.
- For the spatial and full/partial temporal correlation matrices (spatial and temporal NetMats), we started with matrices of size N_mode_ x N_mode_ for each subject. After applying the Fisher r-to-Z transformation, we flattened these matrices by taking the upper diagonal elements. Next, in order to obtain feature matrix from spatial NatMat, i.e. **SNET**_feat_ of size N_subject_ x 500, we used SVD for dimensionality reduction, and reduced the flattened spatial NetMats to 500 features. Similarly, for temporal NetMat feature matrix **TNET**_feat_, we extracted 250 SVD features from each of the full and partial temporal NetMats and concatenated them to obtain a **TNET**_feat_ of size N_subject_ x 500. Partial temporal NetMats were computed based on precision matrices and with a Tikhonov regularisation parameter 0.01.
- Mode amplitudes were used without any transformations; i.e. providing a **H**_feat_ matrix of size N_subject_ x 150.

We conducted *uni-element* predictions based on **P**_feat_, **SNET**_feat_, **TNET**_feat_ and **H**_feat_ separately, and *multi-element* predictions when all these elements were combined. For the multi-element predictions, we utilised pre-prediction univariate feature selections to further reduce from 1650 features to 900. For this purpose, in each cross-validation loop, the top 900 features that showed the highest correlations with the target variable, within the training set, were selected. 5-fold cross-validations were repeated 20 times for prediction of each cognitive score, the 20 repeats are pooled for visualisation using Bland-Altman plots and averaged for statistical comparisons.

#### 3.4.5 Uni-mode prediction

The uni-mode prediction steps are conceptually similar to multi-mode predictions outlined above, with a few exceptions. Firstly, for the dimensionality reduction of the spatial maps we did not need to combine information across modes using bigFLICA. Instead, we used sparse dictionary learning to obtain feature matrices of N_subject_ x 500 for each mode. For spatial and temporal NetMats we take rows of correlation matrices for each subject and do not apply additional dimensionality reduction, thus obtaining 149 features per subject, per mode for each NetMat. For the amplitudes, as there is one value per subject-mode, we simply add a mode’s amplitude to the temporal NetMat feature matrix.

#### 3.4.6 Statistical significance of predictions

We conducted the aforementioned predictions based on both sPROFUMO and ICA-DR, and compared the overall model predication accuracies, as well as the predictions of different model elements in the spatial and temporal domains. In order to estimate p-values, we took note of the fact that cognitive tests included in the study (Table S 1) can be correlated with each other, and as such, a regular paired t-test is not valid because it assumes that the sample covariance is spherical. In order to overcome this problem, we used a generalised least square approach as implemented in MATLAB *lscov* function, which solves a linear regression, A x = B, by finding x that will maximise (B – A x)^T^ V^-1^ (B – A x). Here A is a column of ones, B is the difference of the correlation accuracies for each comparison pair (e.g. ICA-DR SMAPs versus sPROFUMO SMAPs). V is the sample covariance, which we approximate by computing the covariance matrix among the cognitive tests, and projecting it onto the nearest symmetric positive definite matrix. After estimating x and the standard deviation of 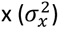, t-values (similar to paired t-test) can be estimated as 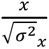, which are then used to estimate the corresponding p-values. Note that for the statistical significance testing, cognitive tests with negative predictions for either of the models in a pairwise comparison are removed as these would act as noise and disguise potentially interesting differences between the two models for the well-predicted tests (Gong et al., 2021).

#### 3.4.7 Canonical Correlation Analysis

Canonical Correlation Analysis (CCA) was used as an add-on test to the prediction results, mainly to disentangle the contribution of spatial versus temporal model elements (section 4.6) in the prediction accuracies of sPROFUMO and ICA-DR. More specifically, it was aimed to test the degree to which sPROFUMO SMAP/TNETs and ICA-DR SMAP/TNETs capture similar sources of subject variability. It was conducted using the following steps:

1. summary features for PFM SMAPs and 500 summary features for ICA-DR SMAPs across all 150 modes were obtained using FLICA, as elaborated in section 3.4.4.
2. summary features for PFM TNETs and 500 summary features for ICA-DR TNETs across all 150 modes were obtained using SVD, as elaborated in section 3.4.4.
3. confounds were regressed out of both PFM and ICA-DR feature matrices, as elaborated in section 3.4.2
4. conducted CCA for four pair-wise comparisons: 1) PFM SMAP vs ICA-DR SMAP; 2) PFM TNET vs ICA-DR TNET; 3) PFM TNET vs ICA-DR SMAP and 4) PFM SMAP vs ICA-DR TNET.
5. CCA would yield a linear transformation of PFM feature matrix (X) and ICA-DR feature matrix (Y) in a way to maximise their correlation; i.e. Y*A=U∼X*B=V, where U and V are the linearly-transformed versions of ICA-DR and PFM feature matrices, respectively.
6. finding correlations between columns of U and V for the top 100 CCA components, we estimated shared variances for each pairwise comparison. We further estimated confidence intervals on correlations using bootstraps.
7. finally tested how many of the CCA components were significantly correlated. For this purpose, we ran permutations: in each iteration, we kept X matrix in step 5 fixed, randomly permuted Y alongside rows (i.e. subjects), and repeated step 6. Across 5000 permutations, we constructed a null distribution for the correlations of the top 100 CCA components. The correlation value corresponding to the top 5% of the null distribution for the *first* CCA component was used as threshold for p-value<0.05 significance level for all the CCA components. This yields a significance threshold that is Family-Wise Error-rate (FWE) corrected for multiple comparisons.

## 4 Results

Results presented here are broadly focused on: firstly, evaluating sPROFUMO’s performance, especially with respect to scaling to big data and reconstruction of a high dimensional set of sPROFUMO modes, or PFMs. Secondly, investigations into the functional and cognitive relevance of these high dimensional modes from resting state data. And, thirdly, a detailed comparison of the model with the ICA & Dual Regression (DR) paradigm, which has been one of the most commonly used methods in RSN research and functional mode discovery.

### 4.1 Simulation set 1 - sPROFUMO vs. PROFUMO

This first simulation set tested how sPROFUMO’s accuracies depend on the choice of the main stochastic parameters, and compared it to PROFUMO. Simulations were designed to mimic real fMRI data (as outlined in 3.1) and comprised two datasets. To mimic big data, each dataset consisted of 500 subjects and 2 recordings per subject. 15 modes were simulated in the brain, which consisted of a mixture of contiguous and distributed modes and were simulated to be temporally correlated and spatially overlapping, where on average 1.3 modes resided per brain voxel. Subject spatial maps (or in short, subject maps) were simulated to have on average of ∼17% spatial misalignment with the group maps (i.e. a given subject mode was on average ∼83% overlapping with the group). Spatial mode layouts were kept constant between subject runs, while mode time courses were allowed to be different, thus resembling the unconstrained nature of the resting-state signals.

As shown in Figure 3, we evaluated the model’s performance by computing correlations between three elements of ground truth and estimated modes: group maps, subject maps and subject time courses. Firstly, we found that sPROFUMO is robust to the choice of batch size, and by decreasing batch sizes from 20% to 5% of the population, model performance was comparable to PROFUMO (Figure 3a). This shows how sPROFUMO can be used to generate the same results as classic PROFUMO while being RAM efficient.

**Figure 3.**
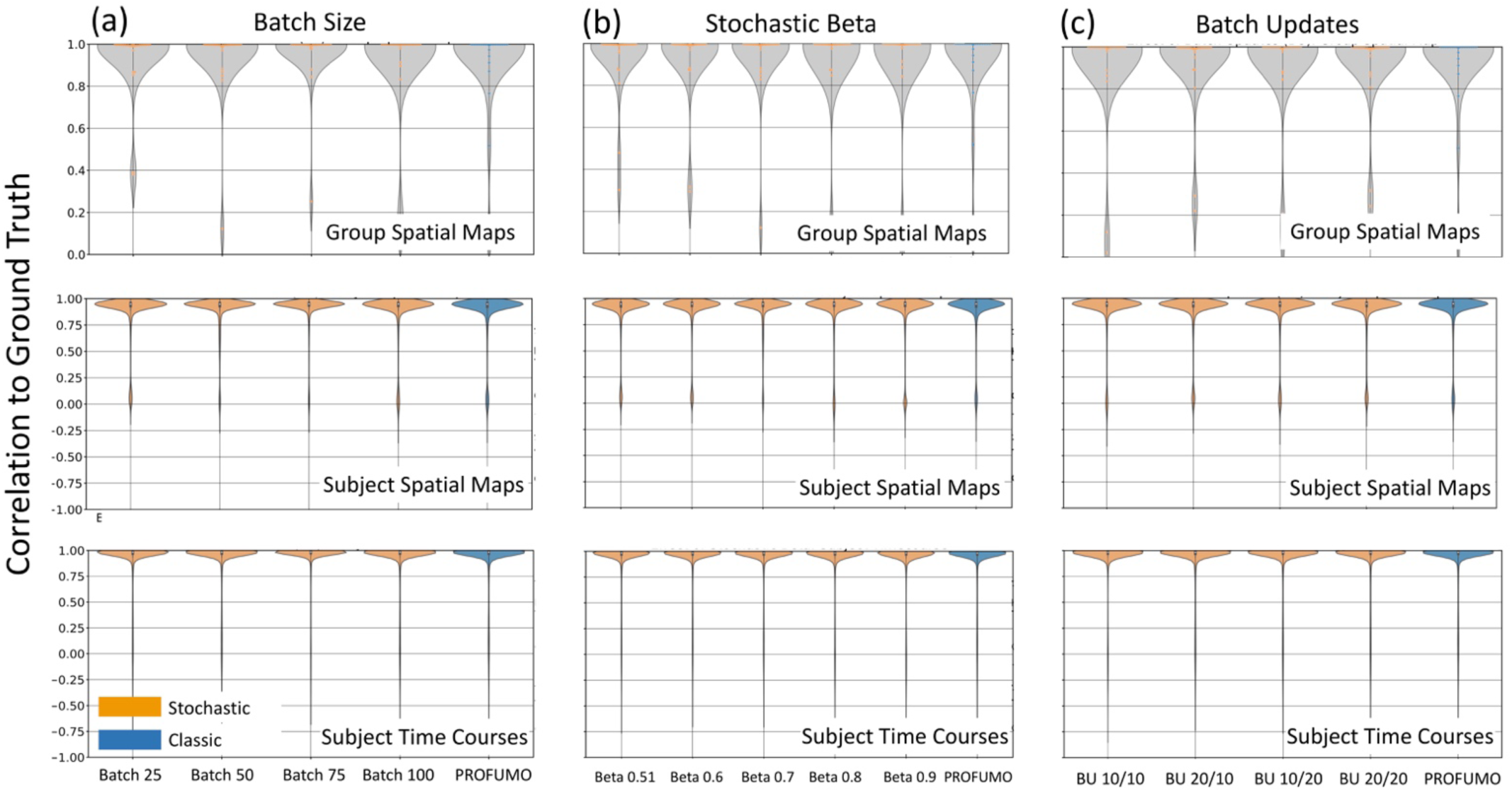
Simulation set 1: illustrating robustness of sPROFUMO (yellow) to the choices of stochastic parameters including batch size, stochastic βand batch updates, and comparison to classic PROFUMO (blue). BU: Batch Updates, BU 20/10 denotes initial/full batch updates.

Secondly, we found that sPROFUMO is robust to the choice of β (Figure 3b). This indicates that in real data application, one can flexibly tweak β values to allow for higher or lower inter-batch variations in the posterior parameters, as would fit data and study requirements. Thirdly, as shown in Figure 3c, we found that the accuracy of the results was robust to the initial and full batch updates (see 2.2.4 for definitions). Thus, in real data this parameter can be flexibly altered to reduce the model’s running time.

This first set of simulations therefore depicts sPROFUMO’s robustness to the choice of specific parameters, thus allowing us to tweak them in real data to obtain more flexible and memory-efficient inference. It is worth noting that for real data applications, it is useful to take data properties such as recording length and noise-level into account when choosing the parameters. For example, for datasets with shorter recordings per subject (e.g. UKB), we recommend using at least 50 subjects per batch to boost within-batch SNR, while for datasets with longer recordings (e.g. HCP), smaller batch sizes can be used. Additionally, for datasets with a small number of subjects and a large number of recording sessions (e.g. Midnight Scan Club (Gordon et al., 2017b)), different recording sessions can be randomised across batches. Moreover, for datasets with excessive noise and artefact levels (e.g. data from babies or participants with movement disorders), using larger batch sizes can be beneficial to increase the effective group-level SNR.

### 4.2 Simulations set 2 - high dimensional RSNs: sPROFUMO vs. ICA

In the next simulation set, we specifically focused on scenarios that are associated with the estimation of a high-dimensional set of modes, and that can be particularly challenging for the matrix factorisation models. In these simulations, we compared sPROFUMO’s performance to ICA-DR. As shown in Figure 4, we focused on three factors; i.e. spatial misalignments across subjects, spatial overlaps among the modes, and mode size, and evaluated each of these factors based on two resting state datasets, each with 500 subjects and 2 runs per subject (see Figure S 1 in Appendix C for examples of simulated and estimated sPROFUMO modes).

**Figure 4.**
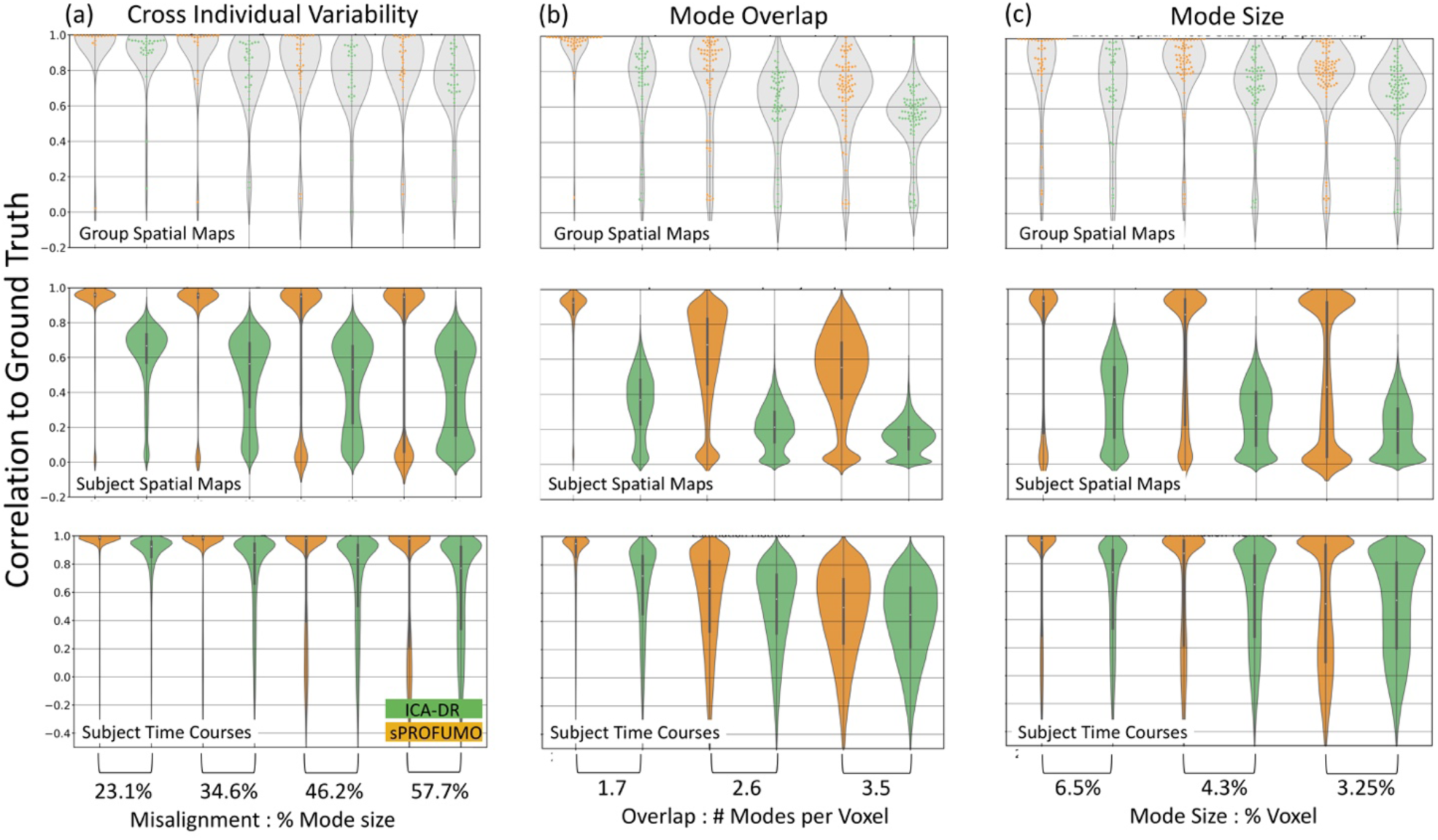
sPROFUMO (yellow) versus spatial ICA (green) in overcoming challenging simulation scenarios for obtaining high-dimensional (i.e. large number of) modes at individual subject level, including: a) cross-individual spatial variability; b) amount of spatial mode overlap; and c) mode size (i.e. smaller modes).

The first scenario tested the hypothesis that in order to obtain a high-dimensional set of modes, we are required to estimate not only the modes that agree across the individuals, but also the modes with larger degrees of cross-subject inconsistencies (e.g. in the presence of spatial misalignments). We simulated 15 modes per subject, and the amount of misalignment (cross-subject) was increased from a moderate 23.1%, where subject maps were on average ∼80% overlapping with the group; to an excessive 57.7%, where subject and group maps were only ∼40% overlapping. In this scenario, simulated modes were set to be non-overlapping at the group-level, and mode sizes were fixed on 8.6% of the brain voxels, on average. As shown in Figure 4a, all the model elements were somewhat affected by the increase in misalignment, but the effect was most pronounced in the subject map estimations. Additionally, the biggest difference between sPROFUMO and ICA-DR was also reflected in the subject map estimations, where e.g. for 57.7% misalignments, sPROFUMO’s average accuracy remained close to 0.9, while ICA-DR’s accuracy dropped to less than 0.5.

In scenario 2 (Figure 4b), we fixed cross-subject misalignments at 300 voxels (i.e. 3% of the brain voxels), and increased the number of modes in the brain from 20 to 30 to 40 while keeping the mode size constant at 8.6% of the brain voxels, on average. Therefore, simulated modes only varied with respect to the spatial overlap between the modes (1.7 to 3.5 modes per voxel on average). We found that while sPROFUMO generally coped well with moderate overlaps (i.e. 1.7 and 2.6 modes per voxel), larger overlaps induced a big impact on both spatial and temporal accuracies of the PFMs. Similar to the previous scenario, we found sPROFUMO’s accuracies to be higher than the ICA-DR. Especially for the extreme case of 3.5 modes per voxel, for the group and subject maps we found: 0.6 ICA vs 0.75 sPROFUMO and 0.15 ICA-DR vs. 0.55 sPROFUMO, respectively. Larger spatial overlap between the modes induces higher spatial correlations among the modes, and as such, this scenario was expected to be particularly challenging for the spatial ICA because it is built around the assumption of spatial mode independence. Somewhat surprisingly, however, we also observed a big impact on sPROFUMO, even though model assumptions were not explicitly violated.

In the third and last scenario (Figure 4c), we held the amount of mode overlap constant, and increased the number of modes in the brain from 20 to 30 to 40, thus obtaining smaller modes, covering 6.5% to 3.25% of the brain voxels. The smaller modes have been frequently reported in high dimensional functional parcellation studies (Dadi et al., 2020; Smith et al., 2015), and are particularly challenging to accurately estimate in single subjects because even small amounts of misalignments (relative to the overall number of the voxels in the brain) can easily surpass the number of the voxels in these modes. Additionally, estimation of these modes is likely to be more affected by the amount of noise in the data and/or dimensionality reduction that is often a part of the preprocessing pipelines. Here we observed that even though sPROFUMO group map and subject time course estimations remained fairly accurate (> 0.8) for the smallest modes, subject map accuracies dropped to ∼0.5. Nevertheless, similar to the previous scenarios, its performance remained superior to that of the ICA-DR by up to 100%.

In summary, these simulations based on three plausible scenarios, provide initial evidence that high dimensional decompositions, particularly at single subject level, are associated with extra challenges for accurate functional mode reconstructions that are not typically seen at lower dimensions. We further showed that the hierarchical modelling in sPROFUMO helps to overcome these challenges often with up to 100% more accuracy than ICA-DR. In Appendix C (and Figure S 2) we show how these results extend to classic PROFUMO and further discuss some of the plausible reasons for when each of these modelling frameworks yield low accuracies. Future studies are expected to investigate additional scenarios and how these findings might generalise to other single subject modelling frameworks.

### 4.3 sPROFUMO on UK Biobank: different types of RSNs identified in a high dimensional soft parcellation

After evaluating sPROFUMO using extensive simulations in the previous sections, we applied the model to the resting state fMRI data from 4999 subjects in UKB and characterised 150 sPROFUMO modes, or PFMs. For this purpose, we set β = 0.6 𝑎𝑛𝑑 τ = 5 for the inference (in Equation 7) and subjects were randomised across batches of size 50. The inference was conducted across 250 batches; thus, each subject was included in a batch 2.5 times on average. Within each batch, subjects were first updated 10 times, keeping the group model constant, in order to bring the subjects into accordance with the group (i.e. *initial* batch updates, see 2.2.4 for details and rationale) and next, the full inference was run 20 times (i.e. *full* batch updates). Therefore, the overall group model was updated 5000 times, and each subject 50 times alongside the group (on average). The model was run on 10 processing cores on a compute node, taking ∼210 hours to complete, and the memory usage peaked at ∼100GB. Model convergence is shown in Figure S 3 (Appendix D), where we measured convergence based on the Free Energy calculated across the full population, as well as the group spatial map and group partial temporal correlation matrix, which represent the two key aspects of the model that are estimated hierarchically.

Figure 5 illustrates the 150 group-level PFMs that consist of a variety of cortical, subcortical and cerebellar modes, providing a high dimensional soft parcellation of the brain. PFMs in this figure are thresholded at 0.5 for illustration purposes. A key distinction between the current results and the previous PROFUMO papers is the reconstruction of a high dimensional set of resting state modes. As shown in Figure 5a, we can broadly categorise these PFMs into four categories: 1) high-SNR distributed RSNs (yellow), 2) lower SNR distributed RSNs (blue), 3) lower SNR parcel-like RSNs (green) and 4) PFMs with physiological or MR acquisition origins (purple).

**Figure 5.**
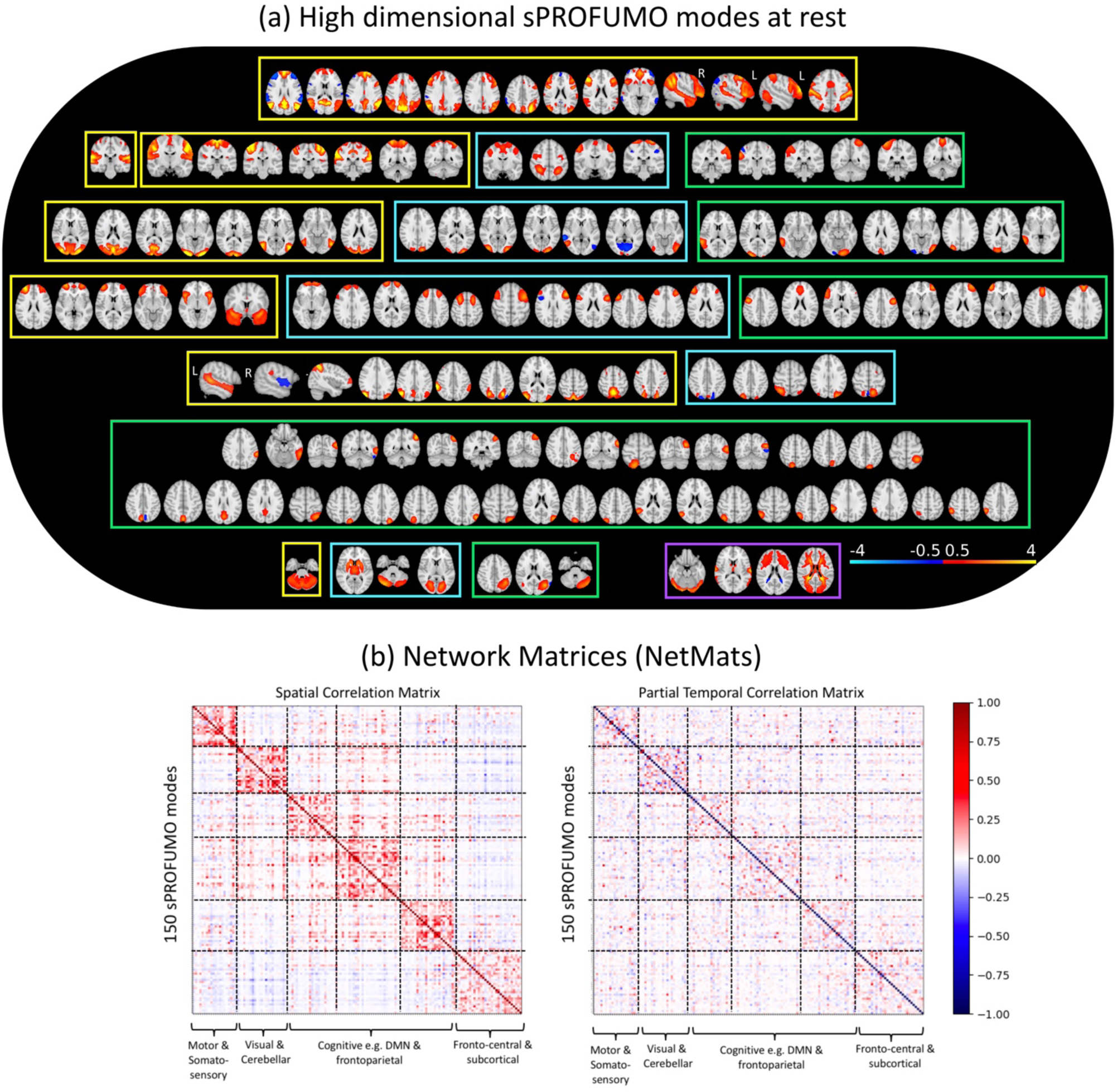
Summary results of applying sPROFUMO to 4999 subjects from UK Biobank and obtaining a high dimensional decomposition with 150 PFMs: a) the reconstructed modes include the large-scale RSNs that are typically found in lower dimensional decompositions, with the addition of less-common RSNs. Four types of PFMs were identified here: high SNR distributed RSNs (category (1) yellow frames), lower SNR distributed RSNs (category (2) blue), contiguous parcel-like RSNs (category (3) green) and PFMs of physiological or acquisition origin (category (4) purple), that altogether provide a high dimensional, soft parcellation of the brain function. Modes shown in this figure are thresholded at an absolute value of 0.5 for illustration purposes. b) Group-average spatial (left) and partial temporal (right) network matrices (NetMats) for sPROFUMO modes. The spatial NetMats cluster into 6 distinct clusters of modes.

Hereafter, we will refer to all the categories as functional modes (or PFMs), but only categories 1-3 are referred to as RSNs. PFMs in category (1) show a close match to the well-known sensorimotor and cognitive RSNs that are typically reported at lower dimensions. PFMs in category (2) appear to represent the variants of sensorimotor and cognitive RSNs that are less frequently found in resting state studies. PFMs in category (3), i.e. the parcel-like modes, were found to spatially overlap with one or several of the distributed RSNs in categories (1) and (2). As will be illustrated later in 4.3.1, we found the signal to noise ratio of the categories (2) and (3) to be lower than the more established RSNs in category (1).

We further characterised the similarities between the PFMs based on two types of correlation-based network matrices (NetMats): partial temporal NetMats and spatial NetMats. As outlined earlier in 2.1, in sPROFUMO (and PROFUMO), modes are simultaneously characterised in both the spatial and temporal domains. In the spatial domain, a mode is described based on its spatial layout (i.e. topography) across the brain voxels and in the temporal domain it is described with time courses, time course amplitudes, and time course partial correlations. The group-level spatial NetMat was computed as cross-mode correlations of the group spatial maps and showed a structured pattern consisting of 6 distinct clusters that were characterised by high within-cluster correlations and low cross-cluster correlations (Figure 5b - LEFT). These clusters were identified using the Louvain algorithm as outlined in (Geerligs et al., 2015) and can be broadly labelled as: somatosensory; visual and cerebellar; variants and subdivisions of DMN; parieto-occipital; fronto-parietal; fronto-central and subcortical. We additionally characterised the group-average temporal NetMat based on partial correlations among the mode time courses, as shown in (Figure 5b - RIGHT). Using the same mode ordering in Figure 5b – LEFT&RIGHT shows that modes with high spatial correlations tend to be generally more temporally correlated, albeit with a notable reduction in the strength of the block structure.

#### 4.3.1 Signal versus noise components

Up to this point, we have shown results based on the “signal” elements of sPROFUMO. However, as outlined in section 2.1, in both spatial and temporal domains, sPROFUMO additionally estimates noise terms. More specifically, every PFM is characterised by six elements: spatial signal and noise, temporal signal and noise, amplitudes and residuals. Figure S 4 in Appendix D illustrates signal and noise components for a set of example resting state PFMs in three random subjects. In the spatial domain (Figure S 4a), it can be seen that the signal element captures the spatial topography of a PFM and its variability across subjects, while the background that is irrelevant to the mode topography is accurately assigned to the noise components. This property is key in obtaining clean subject spatial maps, and can be expected to be particularly useful for datasets with naturally lower SNRs or when less data per subject are available. In the temporal domain (Figure S 4b), the separation of signal and noise components clearly shows how the time courses of example RSNs from categories (1), (2) and (3) from Figure 5 differ from each other, with category (1) showing substantially higher SNR.

#### 4.3.2 Stability of sPROFUMO PFMs across subjects and model runs

We further evaluated how reliably sPROFUMO modes were estimated across and within subjects based on three set of consistency metrics: a) between-run consistency; b) consistency of results when a smaller population is analysed and c) cross-individual robustness of the results. Here we present a brief summary and further details are presented in Appendix E and Figure S 5. Firstly, between-run stability was measured by re-running the model on the same subjects and measuring correlations between the outputs of the two runs, in order to measure stability to stochastic randomisations. We found average consistency of 0.98, 0.98 and 0.94 for the group spatial maps, spatial and temporal NetMats, respectively. For subject-specific PFMs these values were: 0.90, 0.94 and 0.77, with 0.85 consistency for the amplitudes. Secondly, we tested whether the model will produce replicable results if a smaller population is analysed. For this purpose, we re-ran the model on a subset of the original subjects, with 1500 subjects, and compared subject and group results to that of the 4999 population. We found average replicability of 0.96, 0.92 and 0.86 for the group spatial maps, spatial and partial temporal NetMats, respectively. For subject-specific PFMs, these values were: 0.84, 0.72 and 0.62, with 0.89 consistency for the amplitudes. Thirdly, we measured cross-individual robustness of the results based on subject-to-group (S2G) and subject-to-subject (S2S) consistencies. In the absence of a ground truth in the real data, higher S2G and S2S consistencies have been often used as metrics of performance and robustness in single subject modelling (Gordon et al., 2020; Guntupalli et al., 2018). This is particularly due to the fact that we expect the biologically meaningful RSNs to exhibit similar key features across individuals (e.g. right-hand motor network should localise to left motor cortex), and we expect the group model to capture the key elements shared across the population. We found the most-to-least consistent sPROFUMO mode elements to be: spatial NetMats (S2S: 0.79, S2G: 0.89), partial temporal NetMats (S2S: 0.70, S2G: 0.83), spatial maps (S2S: 0.55, S2G: 0.73) and mode amplitudes (S2S: 0.31, S2G: 0.56). It is worth noting that while S2G and S2S consistencies yield useful metrics of results stability, in order to provide more direct evaluation of the model’s ability to capture meaningful subject-specific variability in spatial and/or temporal domains, we complement results from this section with additional metrics from simulations (4.2) and prediction power for cognitive tests (4.6).

### 4.4 Functional relevance of sPROFUMO PFMs at the single-subject level

A closer examination of the PFMs in Figure 6 reveals interesting links with the established functional subdivisions of the brain, not only for the group but also at individual subject level. For example, Figure 6a – RIGHT illustrates four large-scale PFMs distributed over motor cortex and cerebellum that closely correspond to a somatotopy map of the foot, left hand, right hand and face/tongue areas in the brain (Buckner et al., 2011; Haak et al., 2018; Van Den Heuvel and Pol, 2010). Importantly, visual inspection of the example subject PFMs shows rich details of topographical variability across the individuals, whilst the characteristic features of the PFMs, e.g. lateralisation of left versus right-hand motor networks persists. Similarly, Figure 6a – LEFT shows seven PFMs in the occipital lobe that correspond to specific functional sub-regions of the visual cortex (L. Wang et al., 2015).

**Figure 6.**
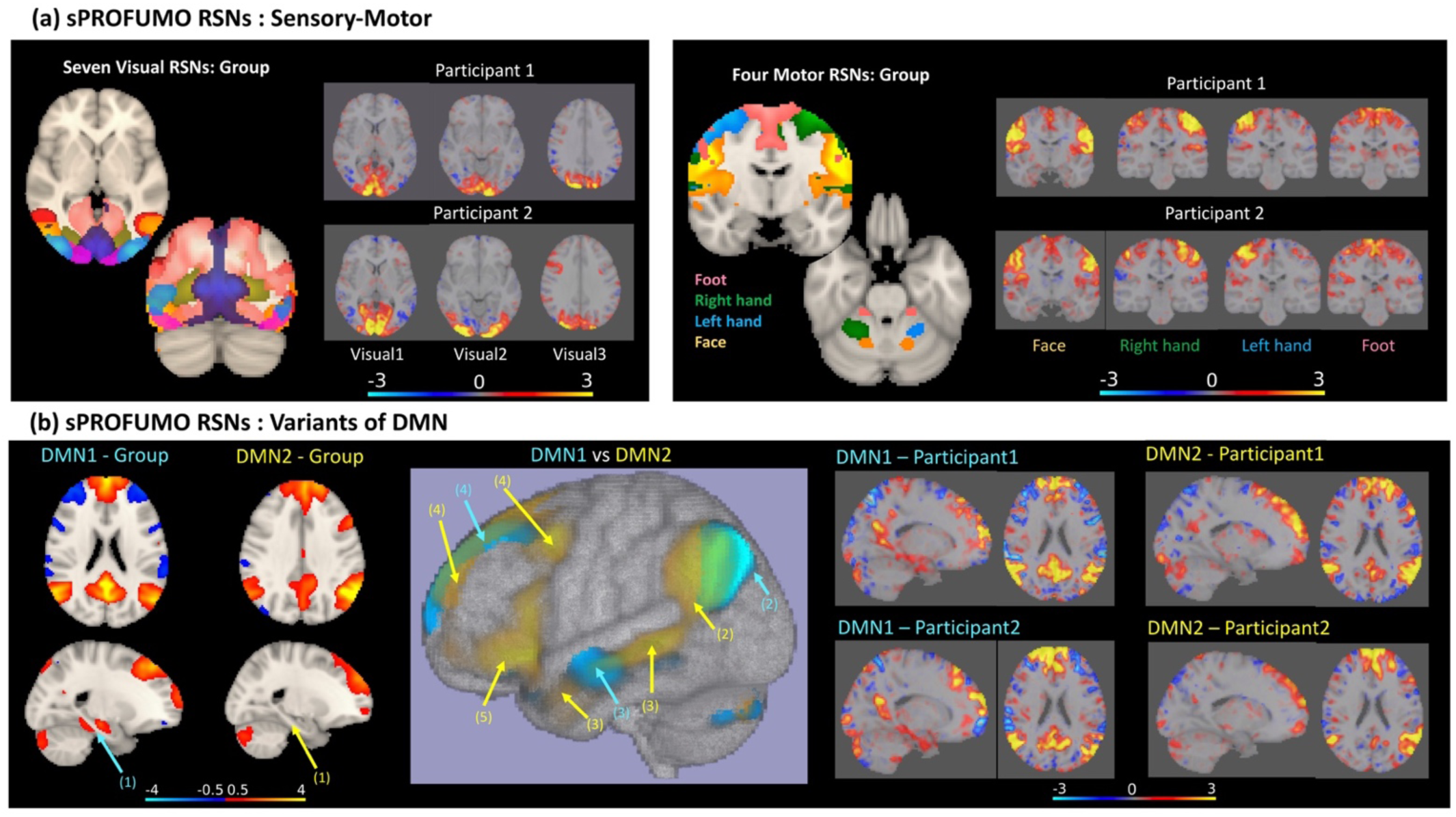
Functional relevance of sPROFUMO modes (i.e. PFMs) in population and individuals illustrated based on: a) examples of visual and motor networks. These PFMs show similarities with the well-known subdivisions of the sensory-motor networks and show rich details of spatial variability across participants; b) two variants of DMN, that are distinct by several factors, marked with arrows, including: 1) a Para-hippocampal node exists in DMN1, but not DMN2; 2) while DMN1 includes posterior IPL and Angular Gyrus, DMN2 includes anterior IPL and Temporoparietal Junction; 3) lateral temporal cortex, including anterior temporal lobes are pronounced in DMN2 but less so in DMN1; 4) while DMN1 includes ventromedial PFC, DMN2 includes dorsomedial PFC; 5) a subnetwork in the inferior frontal cortex exists in DMN2 but not in DMN1. Note that group-level modes shown in this figure are thresholded at an absolute value of 0.5 for illustration purposes but subject-level modes are unthresholded to preserve the details of cross-subject variability.

In addition to the sensorimotor RSNs, we detected multiple variants of the higher-level cognitive networks in single subjects. Figure 6b illustrates two variants of the DMN which is typically characterised by sub-networks in bilateral Inferior Parietal Lobe (IPL), Posterior Cingulate Cortex (PCC), bilateral Temporal Lobes and medial Prefrontal Cortex (PFC). The two DMNs identified here were distinct by several factors (see six marked arrows and explanations in Figure 6b and its caption). Considering these distinctions, DMN1 and DMN2 seem to closely match respectively to the medial temporal and dorsal medial DMN variants that recent studies have reported (Andrews-Hanna et al., 2014; Braga and Buckner, 2017), and which was also reported in the PFMs inferred from HCP data using PROFUMO (Harrison et al., 2020). The important distinction between the results we show here and these previous results is that the characteristic features of the PFMs hold in individual subjects, even when using only six minutes of data per subject: for example, note the presence of the Para-hippocampal subnetwork in DMN1 versus its absence in DMN2. By way of contrast, HCP results used one hour of data per subject, and Braga & Buckner’s analyses used nearly three hours of data per subject.

### 4.5 Distinct behaviours of sPROFUMO and ICA at 150-mode decomposition

We next conducted a detailed comparison of the sPROFUMO PFMs with 150 functional modes estimated using spatial ICA (FSL’s MELODIC) and Dual Regression (DR). Results are shown in Figure 7. Note that group-level modes in this figure are thresholded for illustration purposes (refer to colourbars and figure caption for details).

**Figure 7.**
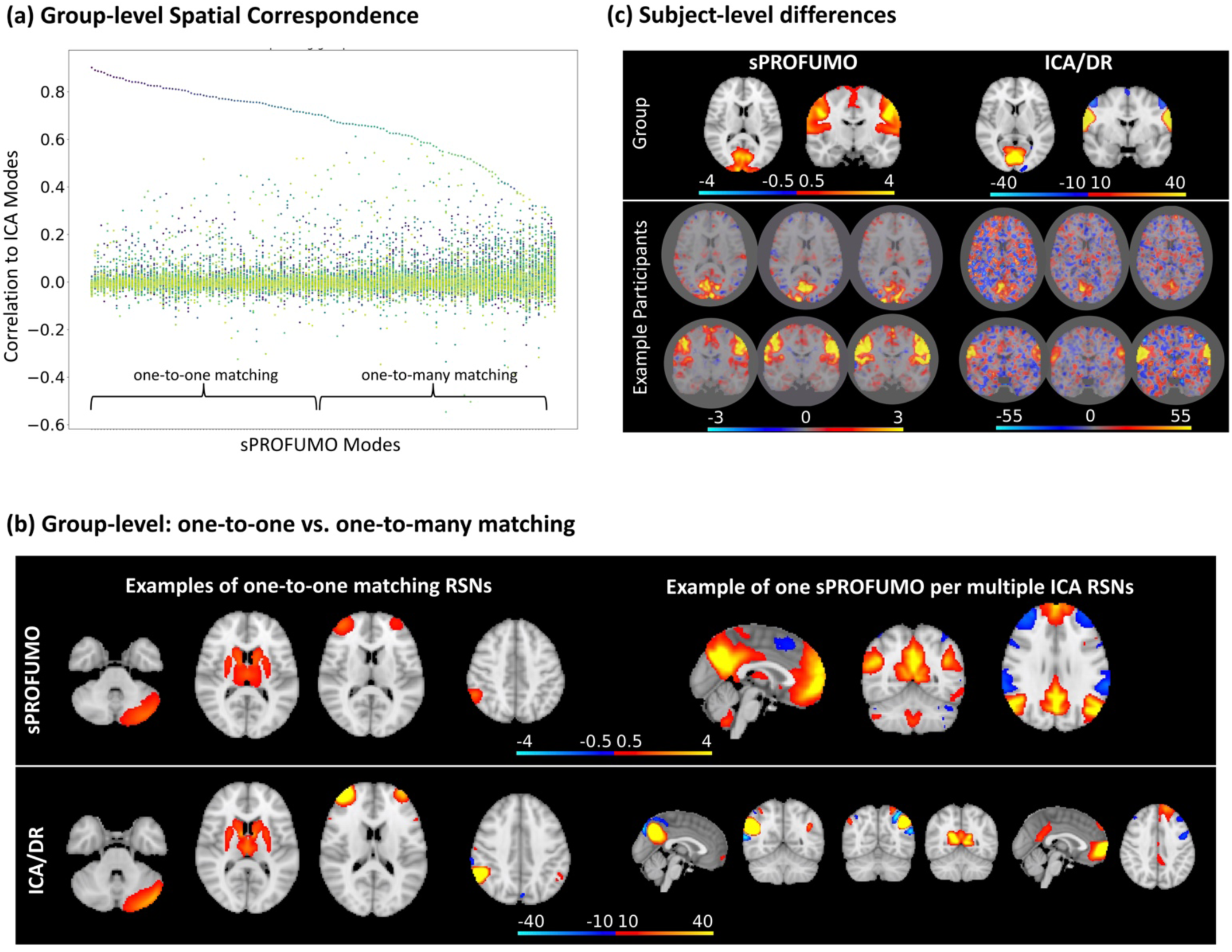
Comparison of 150 sPROFUMO modes from UKB to spatial ICA (group) and ICA-Dual Regression (subjects): a) sPROFUMO and ICA modes that are matched based on spatial correlation across voxels. For the 74 modes labelled as showing one-to-one matching, absolute spatial correlation between each pair was >0.7 and this correlation was at least two standard deviations higher that the next best matching mode. Modes that do not pass this criterion are labelled as having one-to-many matching. b) Left: examples of cerebellar, subcortical, frontal and parietal modes with one-to-one matching; right: example of one DMN mode in sPROFUMO that divided into six subnetworks in ICA; c) example subject-level differences between sPROFUMO and ICA-DR for sensory-motor networks in two UKB participants. Note that group-level modes shown in this figure are thresholded for illustration purposes (refer to colourbars for thresholds) but subject-level modes are unthresholded to preserve the details of cross-subject variability and noise levels in each spatial map.

At the group-level, we paired sPROFUMO and ICA modes based on the spatial correlation across voxels. As shown in Figure 7a, for 74 modes, we found a clear one-to-one correspondence between PFMs and ICs, while for the other half, every PFM corresponded to multiple ICs. A closer examination of the modes revealed that one-to-one correspondence typically occurs for smaller modes (i.e. categories (2) and (3) in Figure 5), such as cerebellar and subcortical RSNs in Figure 7b. In contrast, large-scale PFMs corresponded to multiple ICs as shown e.g. for DMN in Figure 7b. This pattern is likely a by-product of differences in model assumptions. More specifically, ICA assumes spatial independence among the modes and as a result, ICA modes are oftentimes minimally overlapping (Beckmann and Smith, 2004; Bijsterbosch et al., 2019). Consequently, by moving to higher dimensions, ICA is expected to split the large-scale modes into multiple fine-grained modes with small overlaps. On the contrary, sPROFUMO does not impose any such assumption and thus yields a mixture of overlapping large-scale and fine-grained PFMs in high dimensional decomposition. As we will later cover in the Discussion, this can have implications for the interpretability of the sPROFUMO and ICA modes.

In addition to the group-level differences, we also detected interesting differences between sPROFUMO and ICA-DR’s single-subject results. In Figure 7c, we show two visual and motor networks because they provide examples of high SNR modes that are typically reliably estimated in single subjects. As can be noticed, PFM subject maps are less contaminated by the background noise. We showed earlier in 4.3.1 how the Double-Gaussian Mixture Modelling in sPROFUMO accurately assigns uninteresting background to the noise component. Here this is reflected in the histogram of the signal map values across the voxels (Figure S 6 in Appendix F), where for a large number of voxels, signal values are near zero, indicating that noise membership probability has outweighed signal probability. However, such a pattern cannot be systematically captured using ICA-DR and instead, as shown in Figure S 6, it yields one Gaussian distribution across the voxels to account for signal and noise, combined.

### 4.6 Predicting cognitive outcome based on sPROFUMO and ICA-DR

We have so far shown evidence of sPROFUMO’s ability to yield stable and functionally relevant results, and its potential to cope with challenging scenarios for reliable estimation of the high dimensional PFMs. We further examined in detail how the model compares against spatial ICA-DR paradigm. An important next question that might occur to the reader is whether the PFMs inferred by sPROFUMO are indeed capable of accounting for interpretable individual-specific traits; e.g. are these high dimensional PFMs of significant cognitive relevance? We looked to address this question by assessing the sPROFUMO PFMs’ ability to predict a battery of 68 cognitive tests in the UKB data. These tests spanned a range of sensorimotor and higher-level cognitive functions.

First, we computed the prediction accuracies by combining outputs from all the functional modes, which we refer to as multi-mode predictions. For this purpose, as outlined earlier in 3.4.4, spatial maps across all the functional modes where dimensionality-reduced and combined using sparse dictionary learning and bigFLICA (Gong et al., 2021), in order to obtain a set of SMAP features across the subjects. Additionally, spatial and temporal NetMats were dimension-reduced to obtain SNET and TNET features, and mode amplitudes across the subjects were utilised as AMP features. The main purposes of the multi-mode predictions were: a) to determine which of the model elements provide the best predictors, and b) to compare sPROFUMO’s performance to ICA-DR. Bland-Altman plots in Figure 8a show correlation-based prediction accuracies, where values above and below the baseline (at zero) depict higher performances for sPROFUMO and ICA-DR, respectively.

**Figure 8.**
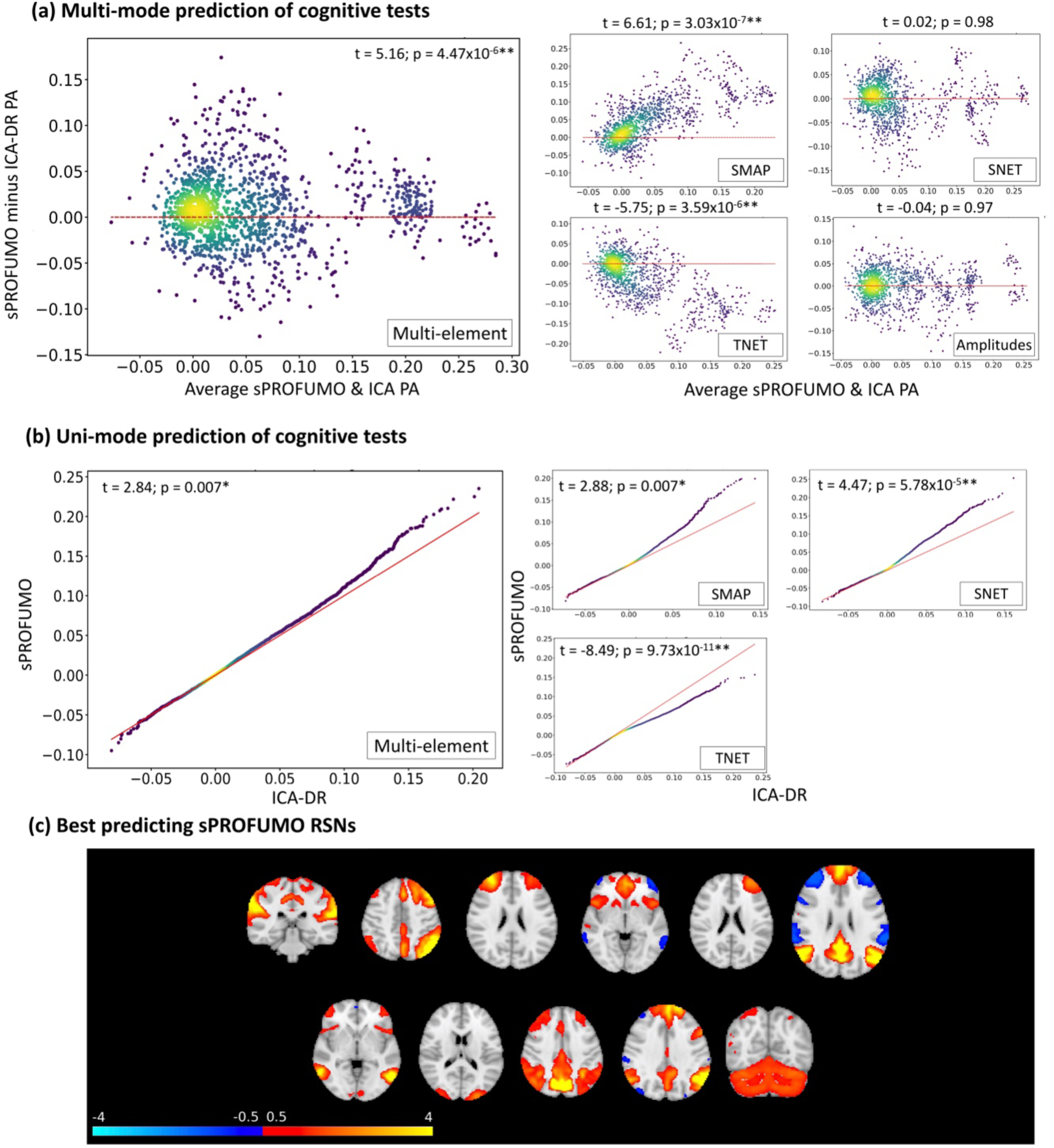
Prediction of cognitive outcome using 150 sPROFUMO modes at rest, and comparison to ICA-DR of the same dimensionality. These results show an overall higher performance for sPROFUMO and reveal a double dissociation between the role of spatial topography versus functional connectivity in prediction accuracies (PA) of the two models. a) Bland-Altman plots for *multi-mode* prediction. 5-fold cross-validations used to estimate prediction accuracies for 68 cognitive tests, each repeated 20 times, providing 20×68 dots per plot; b) density scatter plots of *uni-mode* predictions. 5- fold cross-validations used to estimate prediction accuracies for each of the 150 modes and 68 cognitive tests, providing 150×68 dots per plot. Since PFM and ICA-DR modes are not necessarily one-to-one matching (see FIGURE 7), predictions are sorted and paired based on worst-to-best accuracy; c) examples of the best predicting sPROFUMO RSNs. Red dashed lines in (a) and (b) denote baselines for equal performance.

Two advantages of sPROFUMO that can be noted from these plots are its superior overall (i.e. multi-element) (p-value = 4.47×10^-6^) and SMAP (p-value = 3.03×10^-7^) prediction accuracies. More specifically, the size, locations and spatial arrangement of the functional modes in sPROFUMO were found to provide better predictions compared with ICA-DR. In contrast, TNET from ICA-DR, i.e. the functional connectivity among the modes, outperformed that of sPROFUMO (p-value = 3.59×10^-6^). Apart from this double dissociation of SMAP and TNET, the SNET and AMP elements showed similar performances between the two models. It is worth noting that, as elaborated in 3.4.2, we conducted predictions after a set of imaging confounds were linearly regressed out of both the predictor and target variables. While this can arguably allow us to obtain more interpretable prediction results, we further tested (Figure S 7) that prediction patterns, in particular the double dissociation, is maintained if no deconfounding or unilateral deconfounding of predictor variables are performed.

We next made a closer examination of sPROFUMO’s multi-mode multi-element accuracies for each cognitive test in isolation, as shown in violin plots in Appendix G, Figure S 8, and broadly categorised the 68 tests into three sets: a) sensorimotor metrics that include tests of reaction time and trail making; b) memory and executive functions that include tests of numeric and prospective memory, symbol-digit and pair matching, and matrix pattern completions; c) a range of fluid intelligence tests. We found sPROFUMO to be most successful at predicting tests related to memory and executive functions, where the average prediction accuracies were found to be as high as ∼0.3.

Next, taking a step away from combining outputs of all the functional modes, we used uni-mode predictions, to identify the best predicting modes. As shown in Figure 8c, these included multiple frontal, fronto-parietal and variants of the DMN, in addition to one visual, one auditory, and one cerebellar mode. We found the uni-mode prediction accuracies to be up to 0.15, 0.2, 0.2 and 0.25 for TNET, SNET, SMAP and multi-element, respectively (Figure 8b). Comparison against ICA-DR revealed a generally better performance for sPROFUMO, reflected in SMAP, SNET and multi-element prediction but not the TNET. The superior performance of sPROFUMO for uni-mode multi-element predictions (Figure 8b LEFT) is particularly interesting as it reveals that single PFMs at 150- mode decomposition are of higher cognitive relevance compared to ICA-DR modes of the same dimensionality.

In summary, we found sPROFUMO to successfully predict a range of cognitive tests, and to generally outperform ICA-DR. Notably, we found a double dissociation between the two models, where in sPROFUMO the spatial mode topographies showed the highest contribution in predicting the cognitive outcome, while in ICA-DR, between-mode functional connectivity provided the best predicting element. This dissociation is in-line with previous findings of ICA-DR and PROFUMO (Bijsterbosch et al., 2019, 2018; Harrison et al., 2020; Llera et al., 2019) based on canonical correlations analysis of behavioural and life factors. In this study, we expand those findings using predictions and a sample size that is approximately five times larger.

#### 4.6.1 sPROFUMO versus PROFUMO predictions

The proposed stochastic version of PROFUMO (i.e. sPROFUMO) aims to make the model applicable to big datasets where PROFUMO’s application is not computationally practical. For example, for the 150-mode PFM decomposition based on 4999 UKB subjects, PROFUMO would require ∼7.5TB, which were intractable with our computing resources, while sPROFUMO required ∼100GB. Nevertheless, it is important to show that sPROFUMO’s ability to predict cognitive tests is not worse than PROFUMO. For this purpose, we ran both models on a smaller population consisting of 1500 subjects. We conducted multi-mode predictions using multi-element model summaries, spatial maps, spatial NetMats, temporal NetMats and Amplitudes. Results are shown in Figure S 9, where we found no significant difference between the two models.

### 4.7 Disentangling double dissociation of sPROFUMO spatial map versus ICA-DR temporal NetMats in predictions

Our prediction results revealed that while ICA-DR TNETs outperform sPROFUMO TNETs for prediction of cognitive tests, sPROFUMO SMAPs show superior performance compared with ICA-DR SMAPs. This double dissociation raises a key question, namely, are the two algorithms measuring overlapping sources of subject variability, but projecting them onto different model elements? In order to address this question, we conducted two analyses:

Firstly, we calculated the degree to which sPROFUMO SMAP/TNET share variance with ICA-DR SMAP/TNET across individuals. For this purpose, we extracted multi-mode summaries of each model in spatial versus temporal domains, and obtained N_subject_ x 500 feature matrices for SMAPs and TNETs. We performed Canonical Correlation Analysis (CCA) to measure shared variance in pairwise comparisons, as outlined in 3.4.7. Results are shown in Figure 9. Interestingly, we found the highest shared variance to be between sPROFUMO SMAPs and ICA-DR TNETs, where 77 out of top 100 CCA components where significantly correlated after Family-wise error-rate (FWER) correction for multiple comparisons (see 3.4.7 for details). We also found significant correlations of sPROFUMO TNETs with ICA-DR TNETs (44 CCA components). On the contrary, we only found a few shared components between sPROFUMO SMAPs with ICA-DR SMAPs (1 significant CCA components) and sPROFUMO TNET with ICA-DR SMAP (2 CCA components). Therefore, it is likely that what ICA-DR projects onto functional connectivity matrices is inherently a combination of spatial variability and functional connectivity of RSNs in PFMs.

**Figure 9.**
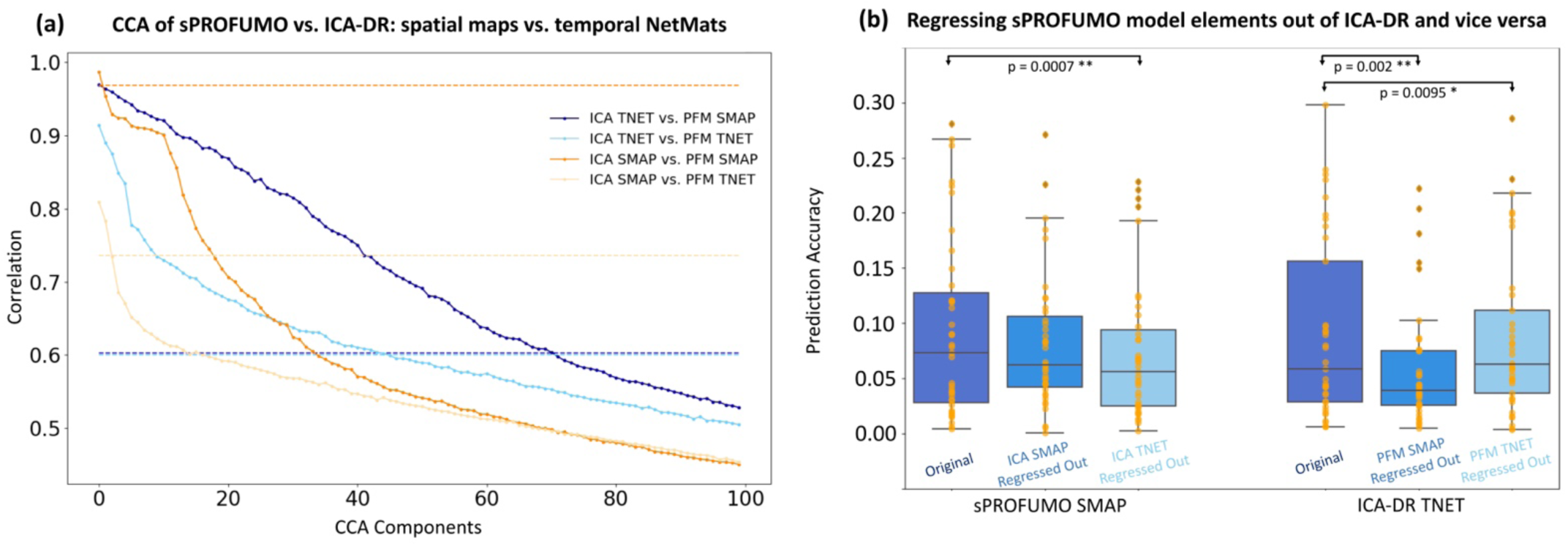
Disentangling sources of subject variability in sPROFUMO versus ICA-DR SMAP and TNETs: a) Canonical Correlation Analysis revealed the highest number of significantly-correlated CCA components to be between sPROFUMO SMAPs and ICA-DR TNETs, followed by shared components between sPROFUMO TNETs and ICA-DR TNETs. Dashed lines show thresholds for significant correlations after Family-wise error rate correction for multiple comparisons using permutations. Pairwise comparisons that are associated with more conservative thresholds showed broader confidence intervals. b) Testing the degree to which sPROFUMO SMAP prediction accuracies will be reduced if ICA-DR SMAP or TNETs are linearly regressed out before running predictions. Testing the degree to which ICA-DR TNET prediction accuracies will be reduced if sPROFUMO SMAP or TNETs are linearly regressed out before running predictions. * denotes p-value < 0.01, and ** denotes p-value < 0.005.

Secondly, considering these significant shared variances between ICA-DR and sPROFUMO model elements, we next tested whether the cognition prediction power of each model will be reduced if the linear projection of the opposite model is regressed out. More specifically, we first focused on sPROFUMO SMAPs, and compared its original prediction accuracies with when ICA-DR SMAPs or TNETs were regressed out (Figure 9b, boxplots on the left). We found that only when the ICA-DR TNETs were regressed out, were the PFM SMAP predictions significantly reduced. Next, we focused on ICA-DR TNETs and compared its original prediction accuracies with when sPROFUMO SMAPs or TNETs were regressed out (Figure 9b, boxplots on the right). This time we found significant reduction of accuracies in both cases. The reduction was particularly evident when sPROFUMO SMAPs were regressed out.

These results altogether confirm that sPROFUMO SMAPs and ICA-DR TNETs measure overlapping sources of subject variability that contribute to prediction of cognitive tests. Importantly, however, while in ICA-DR the sources of cognitive variability seem to be predominantly mapped onto functional RSN connectivity, sPROFUMO appears to divide them between spatial variability and temporal connectivity, with the former playing a more important role. As we will discuss later in Discussion, this can have important implications for how spatial topography and functional connectivity might interplay in subject-specific resting-state network estimations.

### 4.8 Effect of mode dimensionality on group- and subject-level PFMs

Results presented up to this point are based on the decompositions of 150 sPROFUMO PFMs in the brain at rest. Recent studies of subject-specific high dimensional functional parcellations have reported hundreds of functional modes at rest (Dadi et al., 2020; Smith et al., 2015). These findings are often based on the datasets with long recordings in the order of hours e.g. HCP data or combinations of multiple datasets. Given the relatively small number of time points in UKB (i.e. 490 time points corresponding to 6 minutes), here we chose to estimate 150 modes. Nonetheless, this choice remained somewhat arbitrary. We therefore explored the extent to which the key results hold when 100 and 200 PFMs were to be estimated.

First focusing on the group model, we paired group spatial maps based on the spatial correlation across the voxels, as shown in Figure 10a. When comparing 100 and 150 dimensions, we found that there was generally a one-to-one correspondence between the first 100 PFMs, and by increasing to 150, sPROFUMO added new PFMs. We made a similar observation when comparing 150 and 200 PFMs. Modes that started appearing at higher dimensions were spatially correlated with multiple of the existing modes. These new modes were among the lower-ranked modes in sPROFUMO’s matrix factorisation and thus, perhaps unsurprisingly, were found to be typically lower in SNR. We next examined model convergence (Figure S 10 in Appendix H) and found that higher-dimensional decompositions showed a quicker convergence of both group spatial maps and partial temporal NetMats.

**Figure 10.**
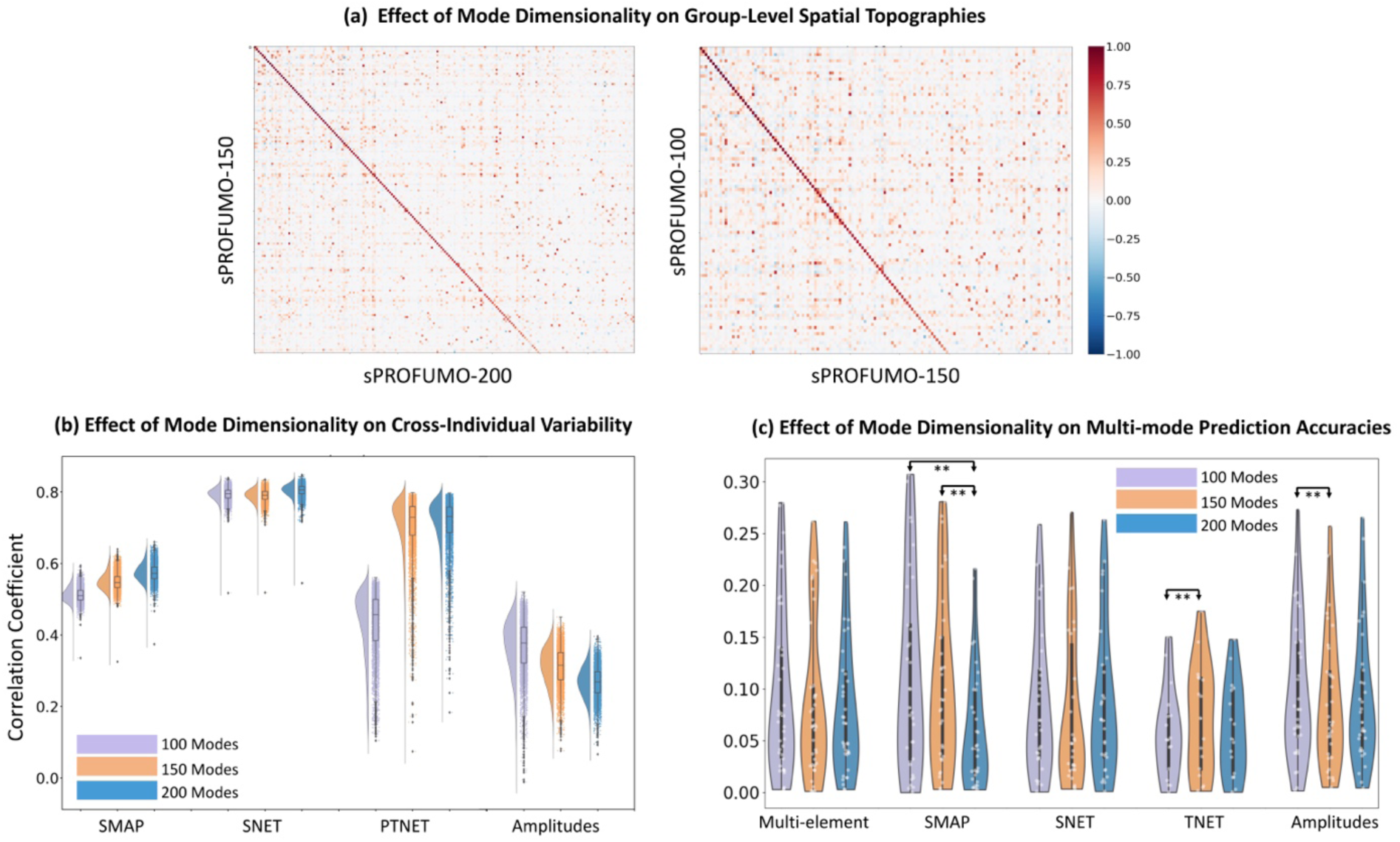
Estimating different number of resting state functional modes using sPROFUMO (comparing 150 to 100 and 200). a) group-level: going to higher dimensions (i.e. from 100 to 150 and 150 to 200) yield new modes that are spatially overlapping with the existing modes; b) subject-level: all model elements except for the amplitudes become more consistent for higher dimensions; c) different dimensionalities show similar multi-mode prediction performances for cognitive tests except for: SMAPs of 100 and 150-mode decompositions are significantly better than 200-mode decomposition, TNETs of 150 are better than 100, and amplitudes of 100 are better than 150. ** denotes p<0.005, refer to the main text for t-value/p-values.

Next, we investigated the effect of increased dimensionality on subject-specific PFMs (Figure 10b). Focusing on cross-subject consistencies, similar to section 4.3.2 and Appendix E, for each model element, we calculated correlations between each subject and all other subjects. Regardless of dimensionality, spatial NetMats were the most consistent model element across the individuals, followed by partial temporal NetMats and spatial maps, and consistencies of these elements further increased with dimensionality. On the contrary, mode amplitudes were the least consistent element and further decreased in consistency by the increased dimensionality. Interestingly, increasing the dimensionality from 100 to 150/200 resulted in more structured partial temporal NetMats with higher off-diagonal values, which in turn resulted in a noticeable increase in cross-subject temporal NetMat consistencies.

We finally compared cognition prediction accuracies of different sPROFUMO dimensionalities (Figure 10c AND Figure S 11). For this purpose, we first looked at the results from multi-mode predictions based on SMAPs, SNETs, TNETs, Amplitudes and multi-element combination. We generally found similar performances for different dimensionalities (Figure 10c), except for three significant differences: SMAPs from 100 (t=4.02; p=0.0003) and 150-mode (t=6.99; p=6.4×10^-8^) decompositions were significantly better than 200-mode decompositions; TNETs from 150-mode were significantly better than 100-mode decompositions (t=3.74; p=0.0018); Amplitudes from 100-mode were significantly better than 150-mode decompositions (t=4.55; p=0.0001). Next, we focused on uni-mode predictions, namely to see if the modes that are found at higher dimensions can outperform the modes that are present at lower dimensions. For this purpose, we focused on 150-mode decompositions. By comparing against 100-mode decompositions, we labelled the top 82 modes that showed a clear one-to-one matching (spatial correlation > 0.75) as primary PFMs, and the remaining 68 as secondary PFMs. We compared multi-element predictions of each secondary PFM to all the primary PFMs and found 7 modes that significantly outperformed at least 25% of the primary PFMs (after FDR correction across 82 x 68 multiple comparisons). Results are shown in Figure S 11, these include 5 parcel-like parietal modes, one frontal and one cerebellar mode.

These results altogether show that the increased dimensionality of the PFMs can add unique information to what is provided by lower dimensions, resulting in more accurate fits to the data and improve consistency. Increasing dimensionality from 100 to 150 (but not from 150 to 200), also showed some improvements for prediction of cognitive tests and revealed new cognitively-relevant fine-grained modes. Therefore, given the relatively short recordings in UKB, 100/150 modes might provide a better choice of dimensionality compared with 200. We will later discuss how our findings on high dimensional PFMs might generalise to other studies in the future.

## 5 Discussion

In this study, we proposed stochastic PROFUMO (sPROFUMO) for inferring functional brain modes in individuals and populations from big fMRI data. We applied the model to resting state data from 4999 UK Biobank (UKB) subjects, in order to obtain high dimensional PFM decompositions of the brain function at rest, and predicted cognitive traits based on these functional modes. The highlights of the proposed model and results are threefold: firstly, sPROFUMO resolves the issue of computational costs that is associated with the application of bidirectional hierarchical models to big fMRI data; e.g. UKB with expected 100,000 subjects. It therefore provides suitable means to leverage the unprecedented richness of population variability that is provided by big data, and explicitly models functional networks for each individual within the epidemiological cohorts. Secondly, using simulations and real data, we illustrated several advantages of sPROFUMO over the established paradigm of ICA Dual Regression. This was particularly reflected in over 100% more accurate estimation of cross-subject variability in the spatial sPROFUMO mode topographies, and yielding more biologically interpretable RSNs at higher dimensions. Thirdly, sPROFUMO enabled us to obtain an unprecedently detailed mapping of the single subject RSNs based on UKB and we showed these RSNs to be predictive of individualistic cognitive traits. The sPROFUMO code and discovered PFMs will be made publicly available in hopes of being useful for future investigations into individualised brain function.

### 5.1 Hierarchical functional mode modelling from big data

An important objective of big neuroimaging data is to provide resources for precision medicine and neuroscience (Bzdok and Yeo, 2017; Ganguli et al., 2018; Quinlan et al., 2020; Sui et al., 2020). Therefore, increasing developments have been made to map the brain function in single subjects, and an important challenge in doing so is to overcome the computational costs of such models (Mejia et al., 2019). Classic PROFUMO (Harrison et al., 2020, 2015) provides an advanced approach for hierarchical modelling of the functional brain networks in population and individuals. The sPROFUMO method proposed in this study builds upon PROFUMO and incorporates two key differences in order to make it suitable for big data application. Firstly, it substantially reduces the computational costs of the model. For example, for a 150-mode PFM decomposition and 4999 UKB subjects as used in this study, classic PROFUMO would typically require ∼7.5TB and for the forthcoming 100,000 subjects in UKB, this value will increase to 150TB of computer RAM, which are intractable. With sPROFUMO, these values are decreased by a factor of 100, such that thousands of subjects can be readily analysed with a modest computing cluster, while still maintaining similar accuracies to that of PROFUMO, as evaluated by extensive simulations and prediction accuracies for cognitive traits.

The stochastic variational inference used in sPROFUMO can enable the model to also accommodate larger degrees of population heterogeneity; i.e. subject deviations from the group. More specifically, in the previous studies, we had observed that PROFUMO requires a mode to be present in a majority of individuals in order to be estimated for the population (Harrison et al., 2020), otherwise, it was found to be typically “switched off” - returned as “missing” for the population as well as all subjects. On the contrary, we found that sPROFUMO was able to recover modes even when they were estimable in only a minority of the subjects. This could be useful in various instances when there are substantial variations in the SNR of the fMRI recordings across the individuals, e.g. in babies or patients, such that the less prominent modes might only be estimable in a subset of subjects. As outlined in 3.2.1, we applied additional data curation strategies to boost the number of estimable modes in individuals.

Another feature of sPROFUMO, which is directly inherited from PROFUMO, and we found to be particularly useful when applying the model to UKB, was its explicit account of signal and noise components at single subject level. UKB, despite the unprecedently large number of subjects that provides excellent group-level SNR, has a relatively small amount of data per subject, where each subject is scanned once and for ∼6 minutes (Miller et al., 2016). This will make it particularly challenging to estimate “clean” modes at single subject level and, thus, further modelling strategies are required to obtain high SNR mode estimation in individuals. Unlike the common approach used in matrix factorisations, where each mode is decomposed into spatial map, time course and residuals (Beckmann et al., 2009; Guo and Tang, 2013; Mejia et al., 2019; Varoquaux et al., 2011), in sPROFUMO, spatial maps and mode time courses are further decomposed into signal and noise components. We showed that (Figure 7 and Figure S 4) this noise modelling can accurately tease apart signal from the background noise and substantially improved the SNR of subject mode estimations, e.g. compared to ICA-DR.

### 5.2 High dimensional single subject RSN decompositions in UK Biobank

We estimated high dimensional sPROFUMO RSNs in population and individuals from UKB. The importance of high dimensional RSN decompositions has come to light in the last few years, as they provide a detailed mapping of the brain function (Dadi et al., 2020; Pervaiz et al., 2020; Smith et al., 2015, 2013b). The fine millimetre spatial resolution of fMRI makes this modality suitable for investigation of the brain function on a voxel-by-voxel basis (Norman et al., 2006; Woolgar et al., 2016). Nonetheless, resting state fMRI connectivity is oftentimes conducted in low-dimensional spaces, e.g. based on 20-50 networks which provides valuable insight into long-distance system-level interactions. Nevertheless, growing evidence suggests that this approach has a limited ability to elucidate the information that is encoded in fine-grained brain topographies (Feilong et al., 2020) or within-mode interactions (Schaefer et al., 2018). The high dimensional modes (100, 150, 200 PFMs) that we identified using sPROFUMO incorporated three types of RSNs: i) high-SNR distributed RSNs, ii) lower SNR distributed RSNs, iii) lower SNR parcel-like RSNs.

Importantly, the high dimensional PFMs demonstrated a unique behaviour whereby the well-known large scale RSNs, e.g. DMN or DAN, that are typically found at lower dimensions were preserved, while new, spatially overlapping, modes were added. This behaviour is distinct from the existing high dimensional functional parcellations (Craddock et al., 2012; Schaefer et al., 2018), and previously reported behaviour in terms of hierarchically organised networks (Doucet et al., 2011; Lee et al., 2012). In particular, this is a fundamentally different behaviour to that which we observed in spatial ICA. Namely, in ICA moving to higher dimensions (e.g. > 100) typically results in the large-scale modes being divided into multiple minimally-overlapping sub-modes that are predominantly confined to one brain region (Kiviniemi et al., 2009; Smith et al., 2015, 2013b). As an example, at 150-mode dimension, we illustrated the DMN in Figure 7, where one PFM spatially overlaps with six different ICs.

The additional RSNs identified with sPROFUMO can be described as the less prominent variants of the well-known RSNs. For example, as illustrated in Figure 5, we found 24 modes in the occipital and occipito-temporal cortex, coinciding with visual networks (L. Wang et al., 2015) while low-dimensional decompositions typically yield less than 10 visual networks (Smith et al., 2012; Thomas Yeo et al., 2011). Our results provide initial evidence for the relevance of these fine-grained modes for predicting cognitive traits. However, future studies are expected to shed more light on details of their functional relevance based on a broader set of phenotypes, ideally based on highly-sampled individuals that can provide sufficient data for high-SNR recovery of fine-grained modes. Some plausible scenarios for the functional relevance of these new RSNs include: a) a subset of these modes that are localised to specific brain regions are likely to be specialised for specific tasks; b) some of these modes might be sub-nodes of the distributed networks such as DMN. These include the modes that show high one-to-one correlations to ICA-DR modes at high dimensions (c.f. Figure 7); c) a subset of these modes, e.g. parcel-like modes in the parietal cortex (Figure 5, 6^th^ and 7^th^ rows), might have occurred due to the limitations of our model for capturing the more complex underlying connectivity patterns e.g. gradients of connectivity, dynamic connectivity, or presence of subpopulations in the data (Lurie et al., 2020; Margulies et al., 2016; Seitzman et al., 2019; Vidaurre et al., 2017).

### 5.3 Predicting cognitive outcome based on functional modes

An important goal of single subject modelling of functional brain networks is to account for individualistic cognitive and clinical traits (Finn & Constable 2016; Gordon *et al*. 2017b; Kong *et al*. 2019). Leveraging the rich source of non-imaging phenotypes in UKB, we used PFMs to predict the outcome of a range of cognitive phenotypes. Based on multi-mode predictions, we found the model to be most successful at predicting tests related to memory and executive functions, where the correlations between the estimated and true test scores were found to be as high as ∼0.3. These prediction accuracies are on par with previous reports based on soft (Pervaiz et al., 2020) and hard parcellations (Kong et al., 2019), as well as the more recent multimodal phenotype discovery methods (Gong et al., 2021). Furthermore, uni-mode predictions unravelled multiple variants of DMN and fronto-parietal attention networks as the best predictors of cognitive phenotypes. This finding can be readily integrated within the previous literature, given the well-established (albeit less well understood) role of DMN and fronto-parietal networks in higher-level cognitive function (Andrews-Hanna et al., 2014; Buckner et al., 2008; Raichle, 2015; Spreng et al., 2010). Interestingly, however, we also found a cerebellar mode as one of the top predicting PFMs, which is in accordance with growing evidence suggesting a key role for the cerebellum in higher-level cognition that goes beyond motor coordination (Schmahmann, 2019; Sokolov et al., 2017; Wagner and Luo, 2020).

Importantly, we found the spatial characteristics of the PFMs, i.e. spatial configuration of the PFMs across the brain voxels as well as spatial correlations among the modes, to provide better predictors of cognitive phenotypes compared to the functional connectivity. While previous research has predominantly investigated links between RSNs and non-imaging phenotypes based on functional connectivity (Dadi et al., 2019; Ng et al., 2016; Pervaiz et al., 2020; Smith et al., 2015), recent studies have highlighted (Bijsterbosch et al., 2018; Kong et al., 2019) that differences in spatial topographies can give rise to trait-like differences. We showed that using sPROFUMO, functional subdivisions of the brain can be clearly reconstructed in individuals (Figure 6), and the prediction results further illustrate that cross-subject variability of these spatial maps is predictive of individualistic traits.

#### 5.3.1 Interpreting the prediction accuracies

In addition to utilising the prediction accuracies as a proxy for the model’s performance, we conducted comparisons between sPROFUMO and ICA-DR in order to shed light on how these results can be interpreted. While we observed a generally higher performance for sPROFUMO, particularly in uni-mode predictions, the most intriguing difference was the increased predictive performance of sPROFUMO’s spatial topography versus ICA-DR’s functional connectivity. This is in-line with recent studies (Bijsterbosch et al., 2018; Llera et al., 2019) and can be explained in light of differences in model assumptions and the way subject-group hierarchies are defined in each framework (Beckmann and Smith, 2004; Harrison et al., 2020; Nickerson et al., 2017).

If we consider the true sources of subject-variability to span both spatial and temporal characteristics of the functional modes, we ideally require a model to: a) identify the variability across the individuals and b) accurately assign the division of labour between spatial versus temporal domains in accounting for that variability. Previous studies have shown that if the spatial variability is not accurately specified, in particular if spatial overlaps between the modes are not accurately accounted for, it can result in a biased, typically over-estimated, functional connectivity (Bijsterbosch et al., 2019, 2018). In other words, true sources of variability that were spatial in nature are likely to be erroneously estimated as functional connectivity and enhance its prediction power. In-line with these findings, based on CCA analysis and predictions, we found that the subject variability that is reflected in ICA-DR TNETs, is mostly captured by sPROFUMO SMAPs and to a lesser extent by sPROFUMO TNETs. Multiple characteristics in the PROFUMO framework are designed to accurately identify spatial versus temporal variability: Firstly, as we showed in simulations (section 4.2), bidirectional hierarchical modelling and inference allows for accurate estimation of spatial mode configurations in single subjects. Secondly, considering that there are no constraints on spatial and/or temporal independence of the PFMs, the model allows the modes to be functionally correlated and spatially overlapping, depending on the evidence provided by the data at hand. Thirdly, and importantly, defining hierarchies on both spatial and temporal characteristics of the modes allows to iteratively fine-tune the parameters in both domains until an optimal balance is found. Therefore, sPROFUMO prediction accuracies can be expected to provide a more interpretable reflection of the true sources of functional mode variability that contribute to cognitive heterogeneity. This can be further tested using other types of phenotypes in the future.

## 6 Data and code availability

Data used in this study is generously provided by UK Biobank and is available upon registration and applying for data access from UK Biobank website: http://www.ukbiobank.ac.uk/register-apply.

Codes used for simulations is available from the following public repository: https://git.fmrib.ox.ac.uk/samh/PFM_Simulations.

Code for sPROFUMO implementation is currently available from the following repository, it will be integrated within PROFUMO and will be made available in an upcoming FSL release: https://git.fmrib.ox.ac.uk/rezvanh/sprofumo_develop.

## 7 Acknowledgements

We would like to thank Dr. Diego Vidaurre for helpful input during early stages of this work. We are thankful to the three anonymous reviewers, who provided valuable feedback, leading to several improvements in this study. We are grateful to UK Biobank for making this invaluable resource available, and to the UK Biobank participants for dedicating their time to make this data possible. SMS is supported by the Wellcome Trust Strategic Award (098369/Z/12/Z) and Collaborative Award (215573/Z/19/Z), and MRC Mental Health Pathfinder grant (MC_PC_17215). MWW’s research is supported by the NIHR Oxford Health Biomedical Research Centre, the Wellcome Trust (098369/Z/12/Z, 106183/Z/14/Z, 215573/Z/19/Z), and the New Therapeutics in Alzheimer’s Diseases (NTAD) study supported by UK MRC and the Dementia Platform UK. SJH was supported by the grant #2017-403 of the Strategic Focal Area “Personalized Health and Related Technologies (PHRT)” of the ETH Domain. SRF is supported by the NIH Human Connectome Project (1U01MH109589–01 and 1U01AG052564–01) and Wellcome Trust (215573/Z/19/Z). The Wellcome Centre for Integrative Neuroimaging is supported by core funding from the Wellcome Trust (203139/Z/16/Z).

This research was funded in whole, or in part, by the Wellcome Trust [Grant numbers 203139/Z/16/Z, 098369/Z/12/Z, 215573/Z/19/Z, 106183/Z/14/Z]. For the purpose of open access, the author has applied a CC BY public copyright license to any Author Accepted Manuscript version arising from this submission.

The computational aspects of this research were partly carried out at Oxford Biomedical Research Computing (BMRC), that is funded by the NIHR Oxford BRC with additional support from the Wellcome Trust Core Award Grant Number 203141/Z/16/Z. The views expressed are those of the author(s) and not necessarily those of the NHS, the NIHR or the Department of Health.

## 8 Authorship contribution statement

Seyedeh-Rezvan Farahibozorg: Conceptualisation, Methodology, Software, Validation, Formal analysis, Investigation, Data Curation, Writing - Original Draft, Writing - Review & Editing, Visualization. Janine D Bijsterbosch: Conceptualisation, Software, Writing - Review & Editing. Weikang Gong: Software. Saad Jbabdi: Conceptualisation; Methodology. Stephen M Smith: Conceptualisation, Methodology, Investigation, Resources, Writing - Review & Editing, Supervision, Project administration, Funding acquisition. Samuel J Harrison: Conceptualisation, Methodology, Software, Investigation, Writing - Review & Editing, Supervision. Mark W Woolrich: Conceptualisation, Methodology, Investigation, Resources, Writing - Review & Editing, Supervision, Project administration, Funding acquisition.

## 10 Appendices

## A Appendix to Introduction

In the Introduction, we focused on three key aspects that distinguish the existing single-subject and hierarchical models of functional brain modes, namely: direction of information flow between population and individuals, hierarchy on spatial topography and/or functional connectivity and scalability to big data. Here we present an extended set of factors, and compare sPROFUMO to some of the existing methods in more detail. The description of these factors is as follows:

- The direction of information flow between population and individuals: whether: a) the group-level estimations are used to regularise subject-specific estimations; or b) subject-specific models are combined to estimate the group model or c) both. We refer to (a) and (b) as Unidirectional (Beckmann et al., 2009; Gordon et al., 2017a; Kong et al., 2019; Mejia et al., 2019; D. Wang et al., 2015) and (c) as Bidirectional models (Abraham et al., 2013; Harrison et al., 2020, 2015; Manning et al., 2018; Shi and Guo, 2016; Varoquaux et al., 2011). Bidirectional models can also be iterative; where the model iterates between group- and/or subject-specific estimations until a convergence point is reached. s/PROFUMO is both bidirectional and iterative, where bidirectionality can be expected to accommodate more cross-subject heterogeneity, and model iterations can be expected to converge on more accurate subject-specific modes through optimisation.
- Explicit versus derivational mode estimations for population or individuals: whether mode estimations are explicitly done for each individual separately (Glasser et al., 2016; Harrison et al., 2020, 2015; Kong et al., 2019; Manning et al., 2018; Mejia et al., 2019; Shi and Guo, 2016; D. Wang et al., 2015) or subject modes are derived as a variant of the group (Abraham et al., 2013; Beckmann et al., 2009; Varoquaux et al., 2011). Furthermore, whether mode estimations are explicitly done for the group, or the group is an average of subjects. Explicit subject-specific modelling in s/PROFUMO can be expected to reduce biases towards population averages, and explicit s/PROFUMO group models can be expected to accommodate more cross-subject variance.
- Defining hierarchy over mode topography or functional connectivity: all hierarchical models attempt at finding a consensus group-level representation, such that subject modes will be comparable. SMAP hierarchy refers to when a model attempts at finding spatial mode layouts that are consistent across individuals (Glasser et al., 2016; Li et al., 2017; Manning et al., 2018; Mejia et al., 2019; Nickerson et al., 2017; Salehi et al., 2017; Shi and Guo, 2016). TNET hierarchy refers to when a model attempts to find between-mode temporal correlation matrices that are consistent across individuals (Chong et al., 2017). Defining hierarchy on both SMAPs and TNETs in s/PROFUMO (introduced in (Harrison et al., 2020)) can be expected to improve the reconstruction of sources of subject variability in spatial versus temporal properties of RSNs.
- Cross-session hierarchy: whether the model allows for an additional hierarchical mode estimation within subjects (i.e. across recording sessions), as incorporated in multi-session hierarchical Bayesian model (Kong et al., 2019). Cross-session hierarchy can be expected to more accurately assign within-subject variations and reduce the possibility of these variations being wrongly attributed to between-subject variations.
- Defining modes as distributed networks (Gordon et al., 2017a; Harrison et al., 2015; Kong et al., 2019; Shi and Guo, 2016; Varoquaux et al., 2011; D. Wang et al., 2015) versus contiguous parcels (Abraham et al., 2013; Glasser et al., 2016; Manning et al., 2018). Both distributed and contiguous parcellations can be beneficial depending on study requirements. As elaborated in 4.3 and 4.8, low-dimensional sPROFUMO PFMs yield distributed functional modes while high-dimensional PFMs yield a mixture of distributed networks and contiguous parcels.
- Hard (Glasser et al., 2016; Gordon et al., 2017a; Kong et al., 2019; D. Wang et al., 2015) versus soft (Beckmann et al., 2009; Harrison et al., 2020, 2015; Manning et al., 2018; Mejia et al., 2019; Shi and Guo, 2016) boundaries between modes. Both hard and soft parcellations can be beneficial, depending on study requirements. sPROFUMO yields soft parcellations.
- Scalability of algorithms to modern big data such as HCP (∼1000 subjects) and UKB (∼100,000 subjects). The latter is one of the main contributions of sPROFUMO, which has not been achieved previously using models that are bidirectional, iterative and incorporate explicit functional mode modelling in population and individuals.

## B Appendix to section 3.2.1: missing modes

In the main section 3.2.1 we discussed the issue of missing modes in PROFUMO framework and briefly outlined five key changes in the data and model that enabled us to largely resolve this issue in this study. Here we elaborate these five factors.

### Reduced group-level missing modes using UKB and sPROFUMO

Here, when applying sPROFUMO to UKB, we found 46/50 and 92/100 non-empty modes at the group-level, which denotes an increase compared to our previous findings (Harrison et al., 2020). A few factors that can explain this observation are: firstly, the large number of subjects in UKB improved the signal-to-noise ratio of the group-level estimations, which can lead to a higher number of group-level modes being reliably reconstructed. Secondly, stochastic inference in sPROFUMO may have allowed for the VB optimisation to jump out of local minima, resulting in an improved minima that seems to allow for a higher degrees of subject variability in the group model. More specifically, in PROFUMO, a mode is typically returned as non-empty only if it is consensually estimable in a majority of the subjects (Harrison et al., 2020). sPROFUMO appeared to show more flexibility around such consensuses, such that modes can be estimated when present in a minority of the subjects too. Finally, in the previous papers we mostly found cortical modes, whereas here a subset of the new modes that we discovered comprised subcortical and cerebellar RSNs. This can potentially be attributed to the type of data, in that here we utilised volumetric fMRI while in the previous papers we used CIFTI (UKB data has not yet been preprocessed into CIFTI form; this is planned for 2021). CIFTI data combines surface-based reconstruction of the cortical areas and volumetric reconstruction of the sub-cortical grey matter. As a result, there are known to be substantial differences in the smoothness of the data measured from cortical and subcortical regions, leading to inhomogeneities in the signal-to-noise ratio. Conversely, volumetric data yields more homogenous signal in different parts of the brain that can potentially help with recovering of the cortical and subcortical modes, alike.

Even though these three factors significantly decreased the number of the group-level missing modes, a number of subject-level missing modes persisted. We applied additional data curation strategies to reduce the remaining missing modes.

### Additional data processing to reduce subject-specific missing modes

Firstly, we applied additional spatial smoothing using FSL tool *fslmaths* with Gaussian smoothing kernel sigma of 2.00, 2.05 and 2.15mm within cortex/white matter, cerebellar and subcortical masks, respectively. The slight difference in smoothing parameters were designed to account for the inherent smoothness differences in resting state signals in these regions. The final estimated smoothness, according to FSL’s *smoothest*, was on average ∼7.2mm FWHM across subjects in all the masks. We observed that this relatively small amount of additional smoothing makes a notable difference to the high dimensional sets of modes obtained from both sPROFUMO and ICA, especially for estimation of single subject spatial maps and reducing missing modes.

As a part of subject-specific preprocessing, PROFUMO by default applies an SVD dimensionality reduction to subject data matrix **D^sr^** in Equation 1 to give **X**^sr^∈ ℝ^N^_v_^xN^_p_, where 𝑁_p_ is the number of top singular vectors. Subsequent matrix factorisations and mode inference are conducted in this dimension-reduced space. For low-dimensional mode decompositions (e.g. ∼40 modes) this dimensionality reduction offers an efficient way of reducing the computational costs while largely preserving the estimation accuracies (provided the number of modes is small compared to the number of timepoints per subject). However, we found that even though the most prominent modes are consistent among the individuals, as we move up to higher dimensions it becomes harder for the SVD to dissociate signal from noise components. Thus, the rank of the modes can arbitrarily change across individuals and more of the group-level modes will be lost in single subjects during SVD reduction. This results in those modes being returned as empty in individuals. In sPROFUMO, considering that only a small fraction of the population is kept in the memory for any given iteration, we are able to work with the full data matrix **D^sr^**. We found this change to greatly reduce the subject-specific missing modes at high dimensions; since, even if a mode does not explain much of the data variance, it can still be recovered in single subjects.

## C Appendix to simulation set 2 (section 4.2)

In the main section 4.2 we presented three simulation scenarios associated with challenges of high-dimensional mode decomposition from resting state fMRI data. Figure S 1 shows examples of simulated ground truth modes and estimated sPROFUMO modes. In the main text, we compared sPROFUMO results to that of ICA and ICA-Dual Regression (ICA-DR). Here in Figure S 2 we show an extended version which also includes classic PROFUMO. Based on these results, in the presence of different degrees of cross-subject variability in the modes, spatial overlaps between the modes and smaller mode sizes, sPROFUMO and PROFUMO show similar performances and they both outperform ICA and ICA-DR. The difference is particularly notable for single subject estimation of the spatial mode topographies where s/PROFUMO tend to yield accuracies that are up to ∼100% superior to that of ICA/-DR.

**Figure S 1.**
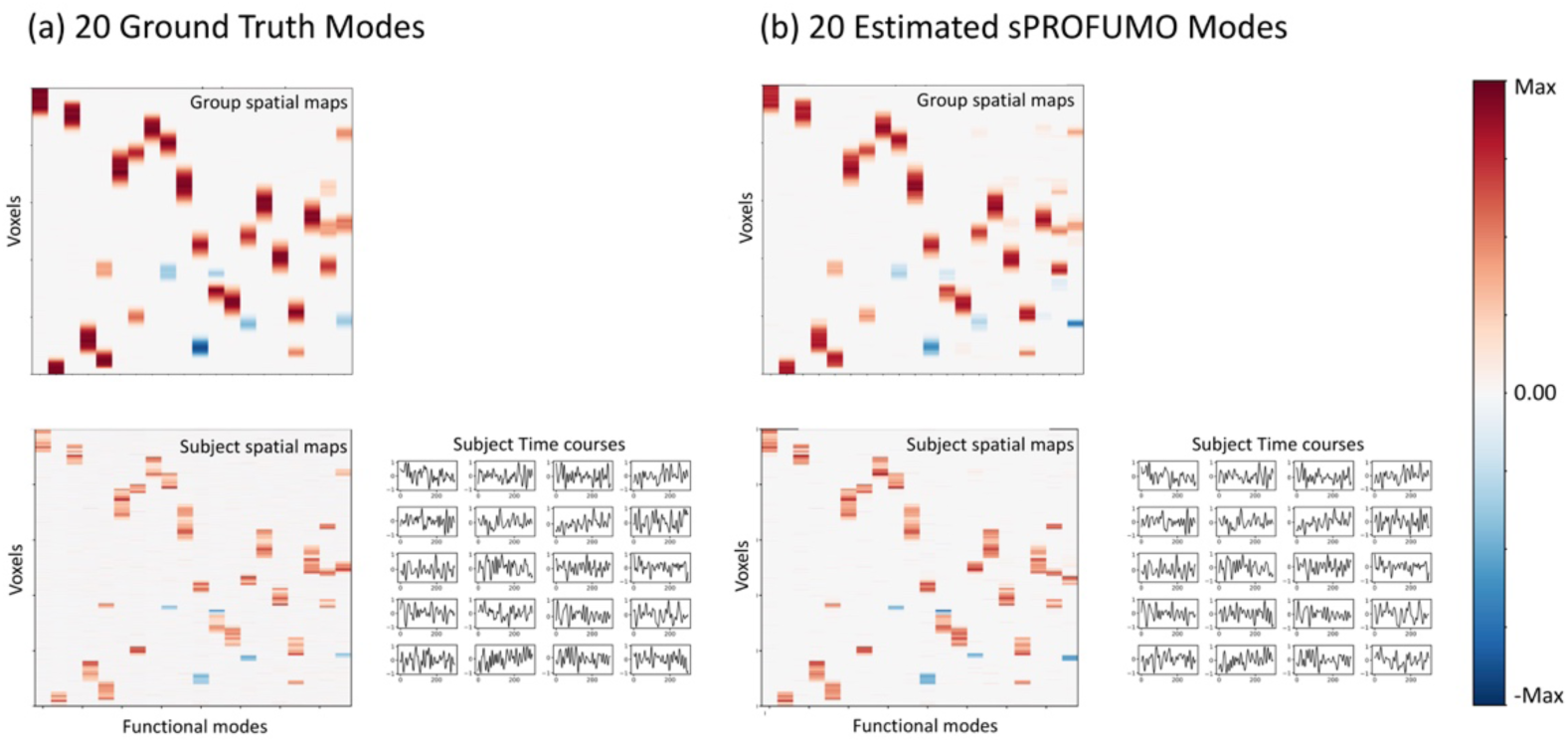
Examples of a) simulated and b) estimated networks in the presence of spatial overlap among the modes and spatial misalignment across subjects. Top: group-level spatial maps. Bottom: subject-level spatial maps and time courses.

### Modes with low accuracies

A closer look at the violin plots in Figure S 2 (and Figure 4) reveals a puzzling behaviour: estimation accuracies of the spatial maps sometimes depict a bimodal distribution where a subset of the modes show accuracies near zero while a majority of the modes are estimated very accurately; e.g. see the distribution where the misalignment is 57.7% in Figure 4a.

A subset of these low accuracy modes can be attributed to when a ground truth mode is missing from the estimations. This is likely to happen at both group- and subject-level, e.g. when a mode is excessively small and/or noisy, such that it cannot be estimated as one of the top modes, and instead is included within the residuals. An additional factor that can give rise to subject-specific missing modes in these simulation scenarios, and consistent with the pattern of results that we observe, is that they occur when a group map is not representative of the entire population. For example, if the amount of misalignment in a subset of subjects is so large that a specific mode has minimal overlap with the group, then it will be difficult for both sPROFUMO/PROFUMO and ICA-DR to estimate those subjects accurately. Arguably, this scenario will also be challenging for other hierarchical algorithms that aim to find one single consensus group model, considering that a key assumption of these models will be violated.

The low accuracy modes are also likely to be false positives; i.e. where a mode that does not exist in the ground truth is detected by the matrix factorisation model. Such false positives might occur when the assumptions of matrix factorisation model (e.g. spatial independence in spatial ICA) are largely violated in the ground truth, or a noise-driven component being erroneously detected as a functional mode.

**Figure S 2.**
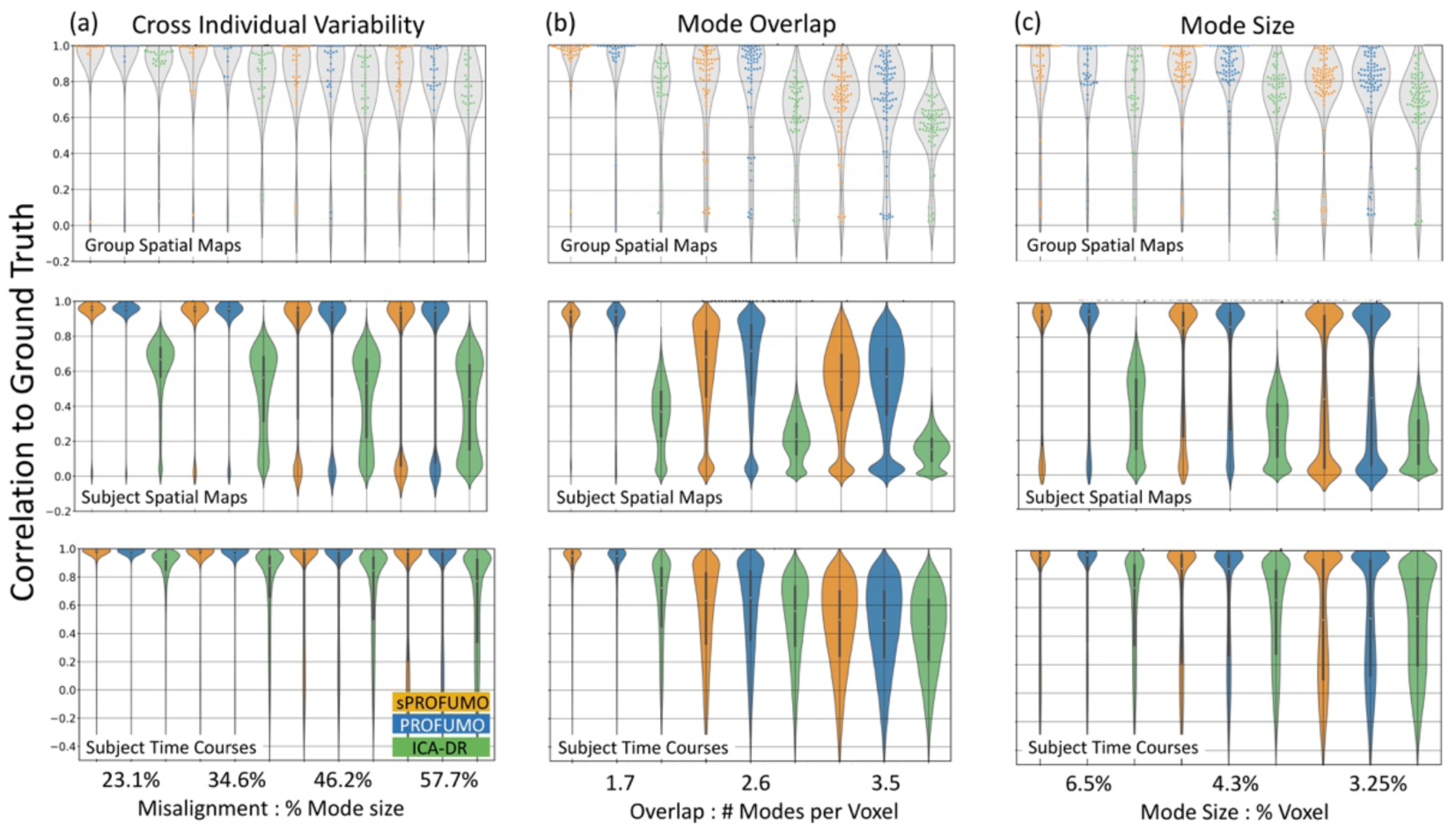
results of simulation set 2 (section 4.2) when including PROFUMO.

## D Appendix to summary of UK Biobank results (section 4.3)

In the main section 4.3 we showed the results of applying sPROFUMO to resting state fMRI data of 4999 subjects from UK Biobank and characterising 150 functional modes. Here we present supplementary figures to that section. Figure S 3 shows model convergence based on full population Free Energy (output from stochastic Variational Bayesian optimisation process) as well as group-level spatial maps and partial temporal NetMats. Figure S 4 is linked to the main section 4.3.1 and shows how explicit decomposition of modes’ spatial and temporal properties into signal and noise components help finding a clean estimation of the subject-specific modes that is minimally contaminated by noise. This also shows how high-SNR large-scale RSNs, low-SNR large-scale RSNs and parcel-like RSNs differ with respect to the noise levels.

**Figure S 3.**
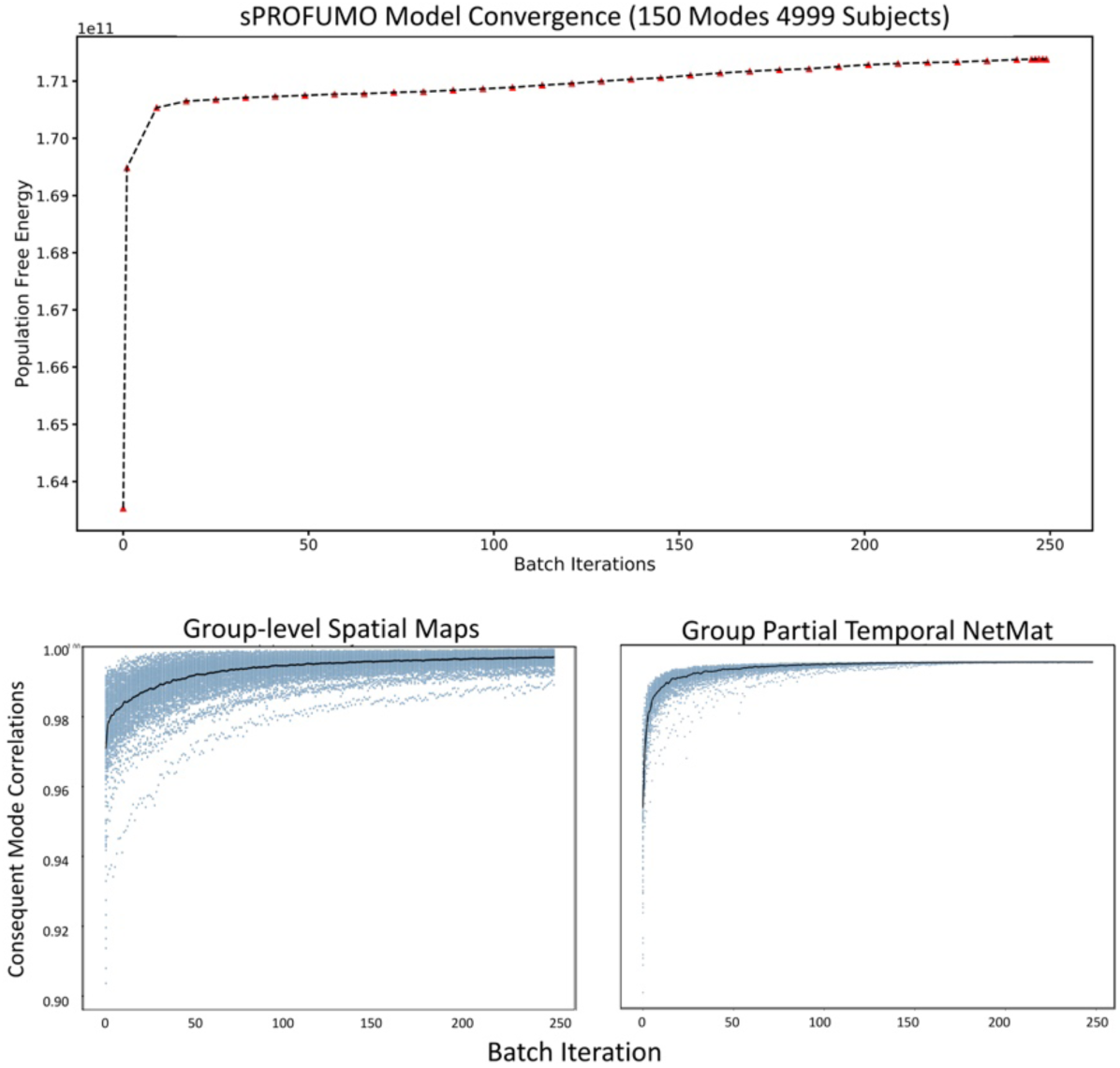
Model convergence on UKB data. Top: Free Energy convergence of the full model; bottom: correlation of group spatial maps and group partial temporal NetMats in each batch iteration (i), with their immediately preceding iteration (i-1) until convergence.

**Figure S 4.**
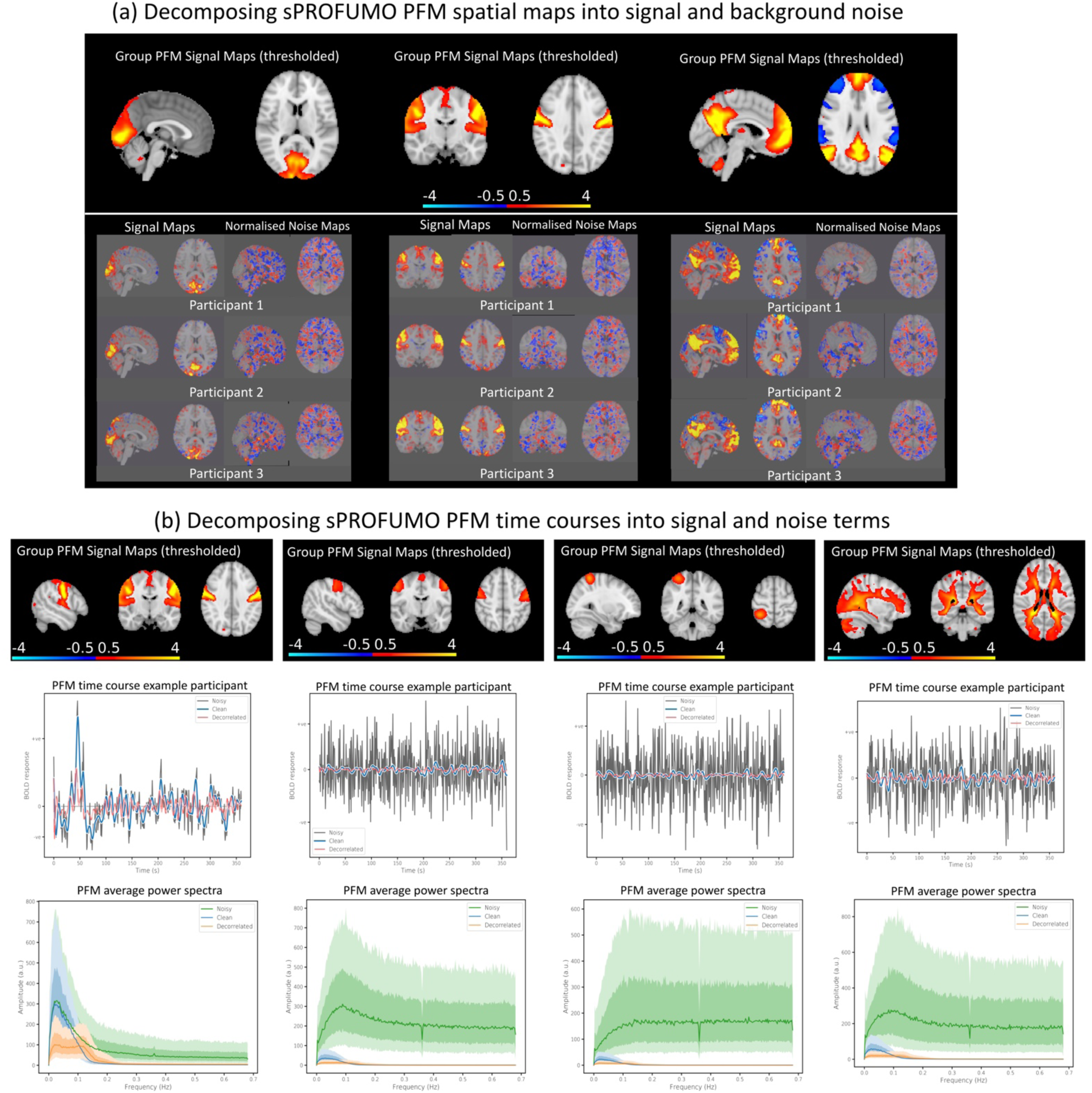
Signal and noise terms in spatial and temporal aspects of sPROFUMO PFMs: a) group-level spatial signal maps (top) and subject-level spatial signal and noise maps for three random participants (IDs: 21126276, 24329265, 24761264). Signal maps in subjects denote the spatial mode topography and noise maps denote the background; b) examples of one PFM per each of the four categories from FIGURE 5: 1) high-SNR distributed (first left), 2) low-SNR distributed (second left) and 3) low-SNR (third left) parcel-like sPROFUMO RSNs in parieto-central cortices as well as 4) a mode with physiological/acquisition origin (right). Note that three timecourses/power spectra are shown in each plot which include: clean time course of the signal component only, decorrelated time course of the signal component after de-convolving with an HRF response function, and noisy time course from combined signal and noise components.

## E Appendix to stability of sPROFUMO PFMs (section 4.3.2)

This section provides a detailed description of the results that were presented in 4.3.2. We evaluated how reliably sPROFUMO modes were estimated across subjects and model runs based on two set of consistency metrics:

Firstly, we tested between-run stability of the sPROFUMO’s results, by re-running the model on the same subjects and measuring correlations between the outputs of the two runs. In stochastic inference, due to the randomisation over the local variables, different runs of a model are prone to yielding different results, even if the inference is conducted based on an identical set of parameters and on the same data. Despite this property, our aim is that the final sPROFUMO output remains stable across multiple runs. To test the stability, we initialised the model based on the same set of initial maps and priors, and re-inferred subject and group PFMs. Results are shown in Figure S 5a where we found average consistency of 0.98, 0.98 and 0.94 for the group spatial maps, spatial and temporal NetMats, respectively. For subject-specific sPROFUMO PFMs these values were: 0.90, 0.94 and 0.77, with 0.85 consistency for the amplitudes.

Secondly, we compared sPROFUMO modes obtained from 1500 subjects with results from 4999 subjects. The 1500 subjects were selected to be a subset of the original 4999 subjects, in order to allow us to test for the replicability of PFMs at both group- and subject-level. While we expect larger population sizes to unravel richer patterns of group and subject variability, this comparison will allow us to test if applying sPROFUMO to smaller populations will produce comparable PFMs with those obtained from larger populations. Results are shown in Figure S 5a, where we found average replicability of 0.96, 0.92 and 0.86 for the group spatial maps, spatial and temporal NetMats, respectively. For subject-specific PFMs, these values were: 0.84, 0.72 and 0.62, with 0.89 consistency for the amplitudes. The lower replicability of temporal NetMats can be traced back to the structure of off-diagonal elements in Figure 5a-RIGHT, where a large number of partial correlations are small and near-zero. These off-diagonal elements can be expected to be affected by noise, thus yielding lower subject-level reproducibility in smaller population.

Thirdly, we measured cross-individual robustness of the results based on subject-to-group (S2G) and subject-to-subject (S2S) consistencies. In the absence of a ground truth in the real data, higher S2G and S2S consistencies are often used as metrics of performance and robustness in single subject modelling (Gordon et al., 2020; Guntupalli et al., 2018). This is due to the fact that, firstly, we expect the biologically meaningful RSNs to exhibit similar key features across individuals (e.g. right-hand motor network should localise to left motor cortex), and we expect the group model to capture the key elements shared across the population. Secondly, the model should ideally be able to remove the effect of measurement noise from single subject mode estimations (e.g. see Figure S 4) which in turn is expected to increase S2G and S2S values. We measured S2G consistencies by finding correlation coefficients of the corresponding mode elements (e.g. spatial maps) between a subject and the group. Similarly, for S2S consistencies, we correlated each subject’s mode elements with all other subjects. Pooling the results in Raincloud plots (Figure S 5a) revealed that S2G consistencies were generally ∼10% higher than S2S consistencies. We further found the most-to-least consistent mode elements to be: spatial NetMats (S2S: 0.79±0.017, S2G: 0.89±0.019), partial temporal NetMats (S2S: 0.70±0.085, S2G: 0.83±0.10), spatial maps (S2S: 0.55±0.024, S2G: 0.73±0.033) and mode amplitudes (S2S: 0.31±0.055, S2G: 0.56±0.093). It is worth noting that while S2G and S2S consistencies yield useful metrics of results stability, they do not inform us of the relationship between estimated and ground truth subject-specific variability in spatial and/or temporal domains. Therefore, we complement results from this section with additional metrics from simulations (4.2) and prediction power for cognitive tests (4.6) to illustrate model’s ability to accurately and meaningfully capture cross-subject variations.

**Figure S 5.**
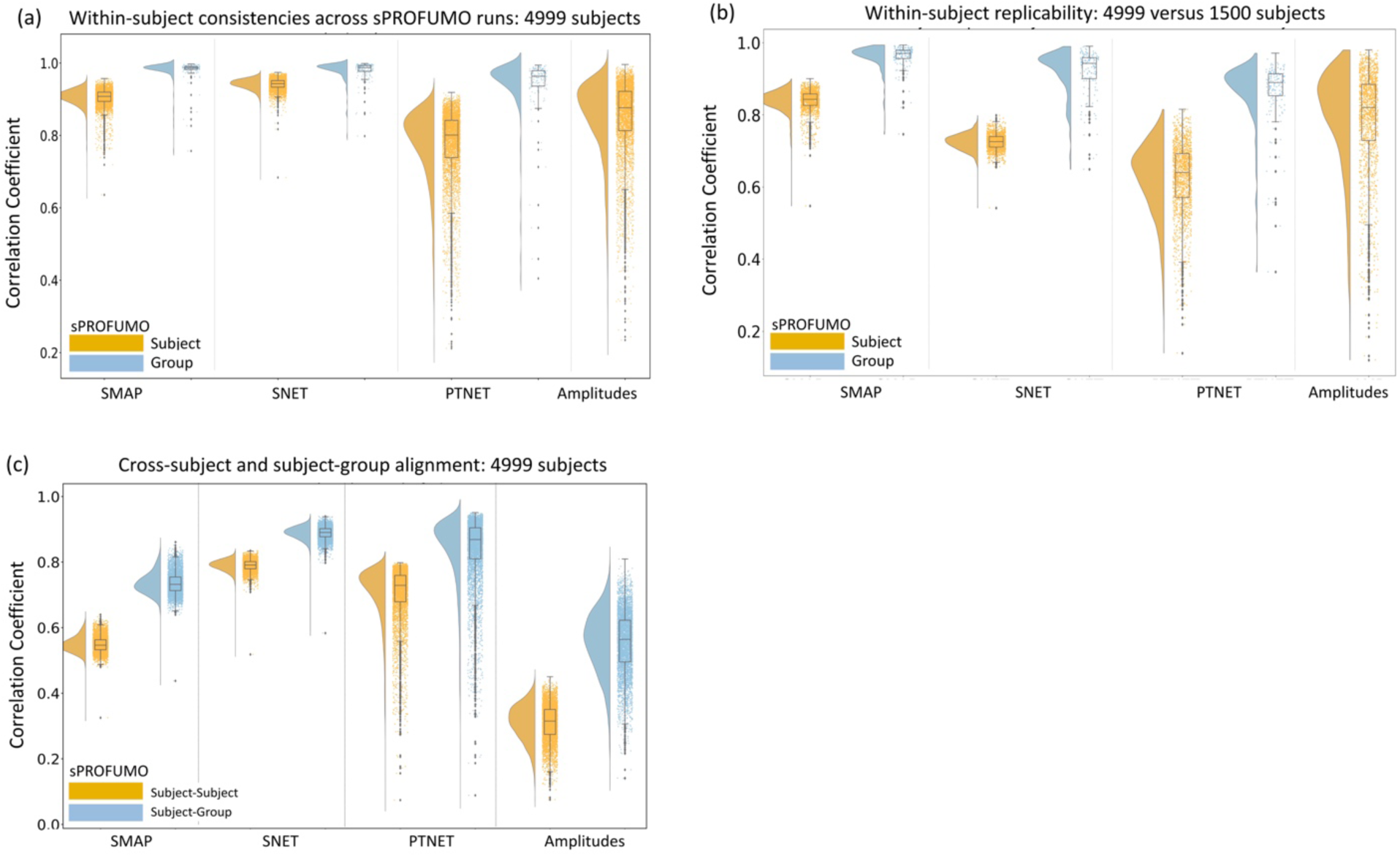
Stability of sPROFUMO modes: a) consistency of group- and subject-level PFMs across two sPROFUMO runs; b) consistency of group- and subject-level PFMs when running sPROFUMO on two population sizes, 1500 and 4999 subjects; c) subject-to-group and subject-to-subject consistency of PFMs within a single model run. Consistencies are measured based on different model elements including spatial maps (SMAP), spatial and partial temporal NetMats (SNET, PTNET), and Amplitudes. Note that we do not include a Raincloud plot (Allen et al., 2019) for the group-level amplitudes in panel (a) because each mode has one group-average amplitude thus yielding 150 values per model run. The correlation between these amplitudes is therefore just a single number.

## F Appendix to comparison of sPROFUMO and ICA-DR (section 4.5)

In the main section 4.5 we showed how high-dimensional sPROFUMO decomposition of 150 modes compared to that of ICA and ICA-DR. Figure S 6 is supplement to FIGURE 7 and shows how distribution of subject spatial maps in different brain voxels differs between sPROFUMO spatial signal element and ICA-DR.

**Figure S 6.**
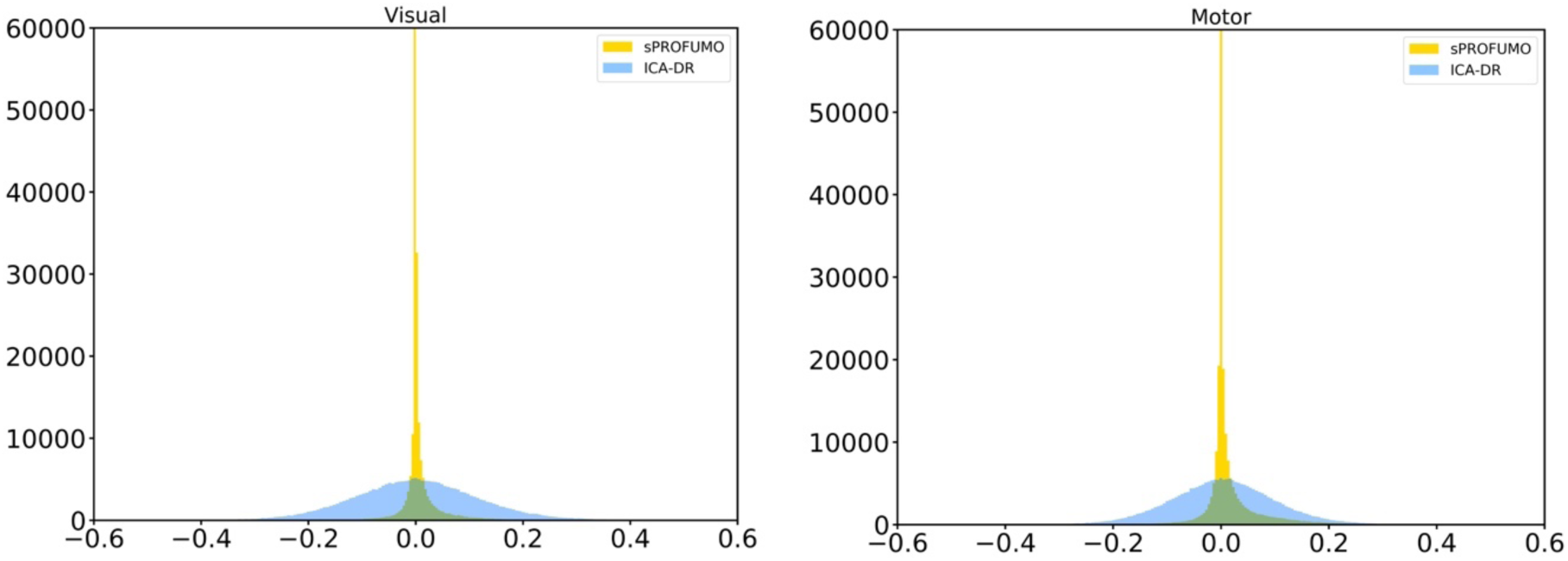
Supplement to Figure 7: example histograms of sPROFUMO subject spatial maps (signal element) and ICA-DR spatial maps across voxels for participant 21126276. Left: visual modes in Figure 7, Right: motor modes in Figure 7.

## G Appendix to prediction results (section 4.6)

In the main section 4.6 we showed results of using 150 sPROFUMO modes from resting state fMRI from 4999 UK Biobank subjects to predict cognitive outcome.

Figure S 8 shows multi-mode multi-element prediction accuracies for every cognitive test separately (refer to the main text for additional explanations). Table S 1 shows the names of cognitive tests included in predictions and full details of each test are available in UKB website: https://biobank.ctsu.ox.ac.uk/crystal/label.cgi?id=100026. Figure S 9 compares sPROFUMO’s multi-mode prediction accuracies to PROFUMO for 1500 subjects and 150-mode decomposition.

**Figure S 7.**
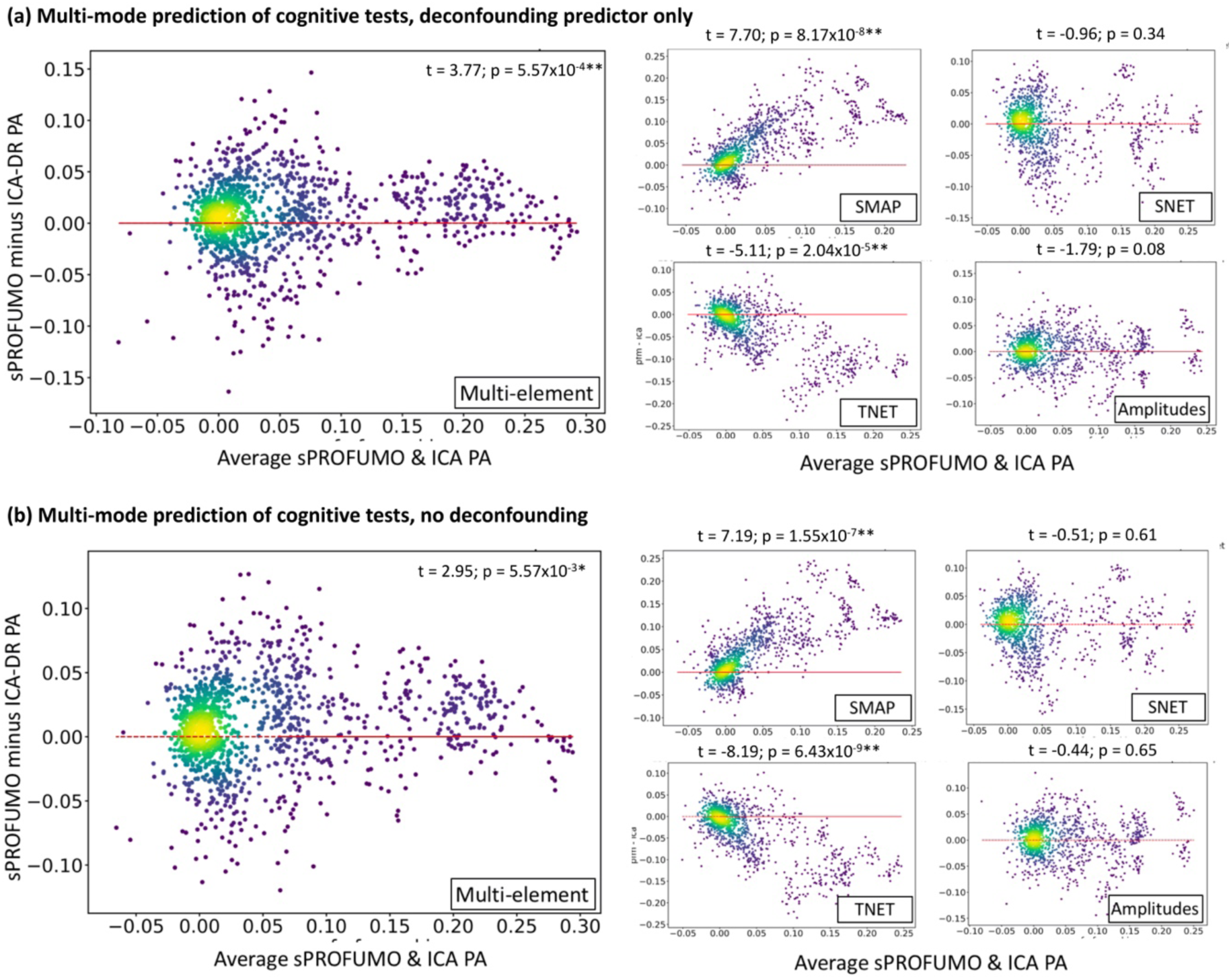
Multi-element prediction results based on different deconfounding strategies: a) deconfounding predictor variable only; b) no deconfounding. Comparison to Figure 8a shows that results and conclusions are maintained regardless of the choice of deconfounding.

**Figure S 8.**
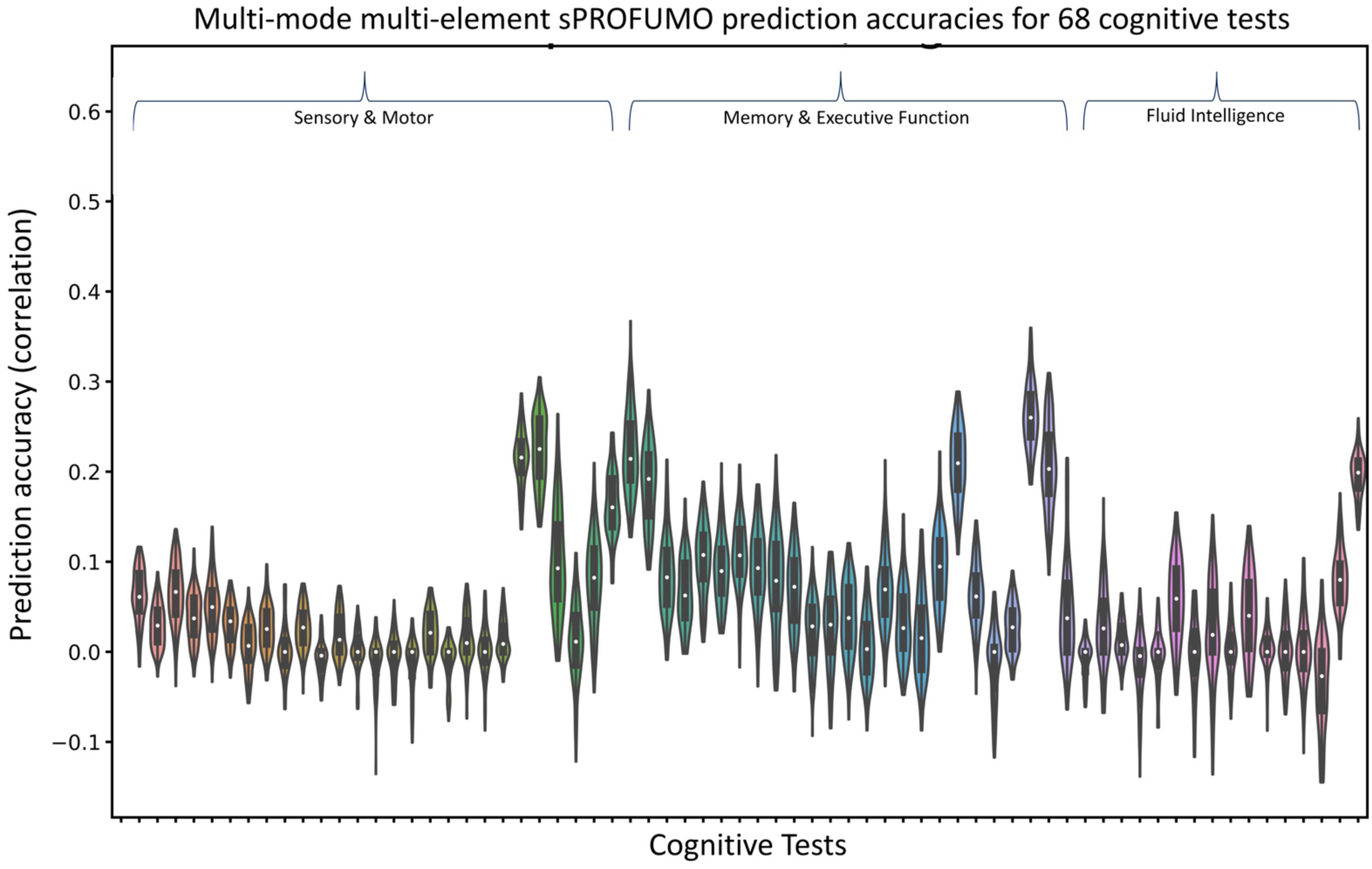
Accuracies of sPROFUMO’s multi-mode multi-element predictions for each of 68 cognitive tests. 5-fold cross-validations are repeated 20 times to obtain distributions for violin plots. The model is generally best at predicting higher cognitive functions such as memory and executive function followed by sensorimotor function and fluid intelligence scores. Best-predicted tests include: Mean time to correctly identify matches (0.0), Mean time to correctly identify matches (2.0), Duration to complete alphanumeric path (trail #2) (2.0), Duration to complete numeric path (trail #1) (0.0), Duration to complete alphanumeric path (trail #2) (0.0), Time elapsed [in numeric memory test] (2.0), Time elapsed [in numeric memory test] (2.1), Maximum digits remembered correctly (2.0), Number of puzzles correctly solved (2.0), Number of symbol digit matches made correctly (0.0), Number of symbol digit matches made correctly (2.0), Fluid intelligence score (2.0).

**Figure S 9.**
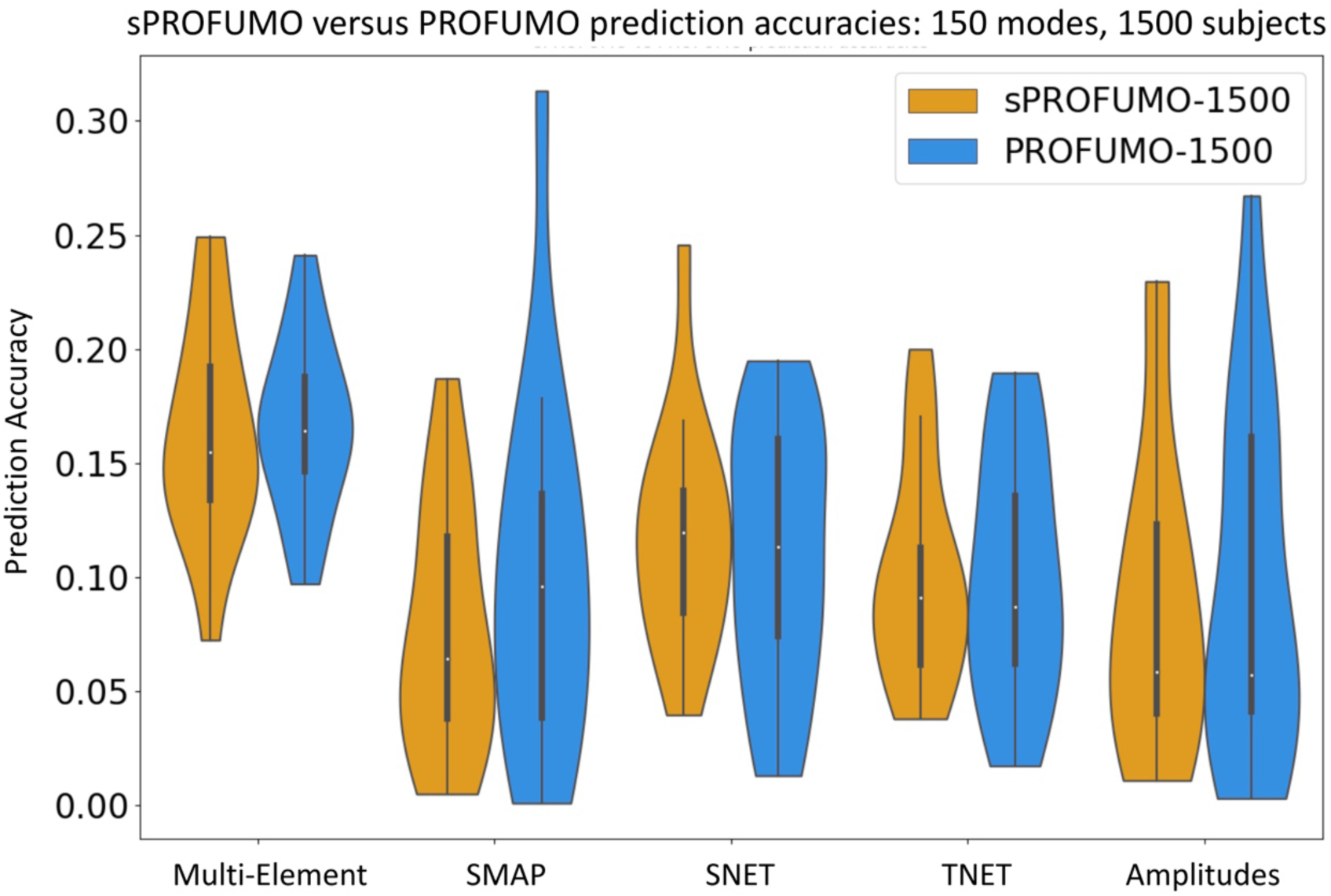
Comparing prediction accuracies of PROFUMO and sPROFUMO: multi-mode predictions used to compare model performances based on 150-mode decompositions of 1500 subjects. Pairwise comparisons based on weighted paired t-tests revealed no significant differences between the two models (all p-values > 0.3). SMAP: Spatial Maps; SNET: Spatial NetMats, TNET: Temporal NetMats.

**Table S 1.** 68 cognitive tests used for prediction.

| Cognitive tests |  |  |
| --- | --- | --- |
| 1- No. incorrect matches (0.1) | 31- Digits correctly (2.1) | 60- Prospective memory (2.0) |
| 2- No. incorrect matches (0.2) | 32- Digits correctly (2.2) | 61- Mn time identify matches (0.0) |
| 3- No. incorrect matches (2.1) | 33- Digits correctly (2.3) | 62- Mn time identify matches (2.0) |
| 4- No. incorrect matches (2.2) | 34- Digits correctly (2.4) | 63- No. incorrect matches (0.0) |
| 5- No. incorrect matches (2.3) | 35- Digits correctly (2.5) | 64- Dur num. path (trl #1) (0.0) |
| 6- No. times snap-button (0.0) | 36- Digits correctly (2.6) | 65- Dur alphanum path (trl #2) (0.0) |
| 7- No. times snap-button (0.1) | 37- Digits correctly (2.7) | 66- No. symbol digit matches (0.0) |
| 8- No. times snap-button (0.2) | 38- Max digits correctly (2.0) | 67- No. symbol digit matches (2.0) |
| 9- No. times snap-button (0.3) | 39- FI1 : numeric add. (0.0) |  |
| 10- No. times snap-button (0.4) | 40- FI1 : numeric add. (2.0) |  |
| 11- No. times snap-button (0.10) | 41- FI3 : word interpol. (0.0) |  |
| 12- No. times snap-button (0.11) | 42- FI3 : word interpol. (2.0) |  |
| 13- No. times snap-button (2.0) | 43- FI4 : posit. arithmetic (0.0) |  |
| 14- No. times snap-button (2.1) | 44- FI4 : posit. arithmetic (2.0) |  |
| 15- No. times snap-button (2.2) | 44- FI5 : fam. rel. calc. (0.0) |  |
| 16- No. times snap-button (2.3) | 45- FI5 : fam. rel. calc. (2.0) |  |
| 17- No. times snap-button (2.4) | 46- FI6 : condit. arith. (0.0) |  |
| 18- No. times snap-button (2.5) | 47- FI6 : condit. arith. (2.0) |  |
| 19- No. times snap-button (2.7) | 48- FI7 : synonym (0.0) |  |
| 20- No. times snap-button (2.10) | 49- FI7 : synonym (2.0) |  |
| 21- No. times snap-button (2.11) | 50- FI8 : chained arithmetic (2.0) |  |
| 22- Time elapsed (2.0) | 51- FI9 : concept interpol. (2.0) |  |
| 23- Time elapsed (2.1) | 52- FI10 : arith. seq. recog. (2.0) |  |
| 24- Time elapsed (2.2) | 53- Dur. num. path (trl #1) (2.0) |  |
| 25- Time elapsed (2.3) | 54- Total err num. path (trl #1) (2.0) |  |
| 26- Time elapsed (2.4) | 55- Dur alphanum. path (trl #2) (2.0) |  |
| 27- Time elapsed (2.5) | 56- No. puzzles correct (2.0) |  |
| 28- Time elapsed (2.6) | 57- Fluid intelligence (0.0) |  |
| 29- Time elapsed (2.7) | 58- Fluid intelligence (2.0) |  |
| 30- Digits correctly (2.0) | 59- Prospective memory (0.0) |  |

## H Appendix to effect of mode dimensionality (section 4.8)

In section 4.8 we showed the effect of mode dimensionality on sPROFUMO results, comparing 150 modes with 100 and 200. Figure S 10 shows the convergence rates of the group spatial maps and partial temporal NetMats based on correlations between the group model obtained in each model iteration with the immediately preceding group model.

**Figure S 10.**
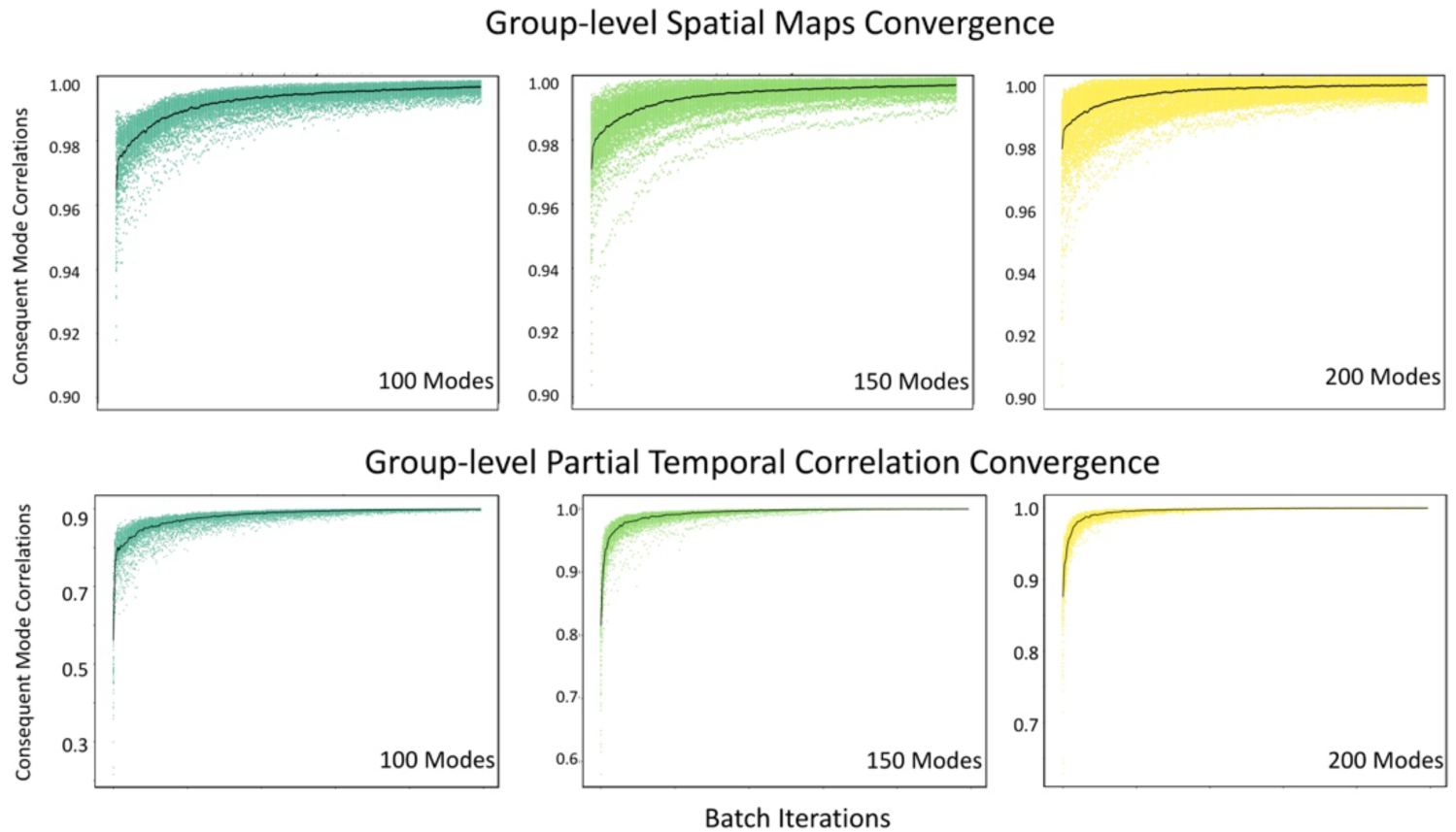
Convergence rates of different dimensions of sPROFUMO modes at rest.

**Figure S 11.**
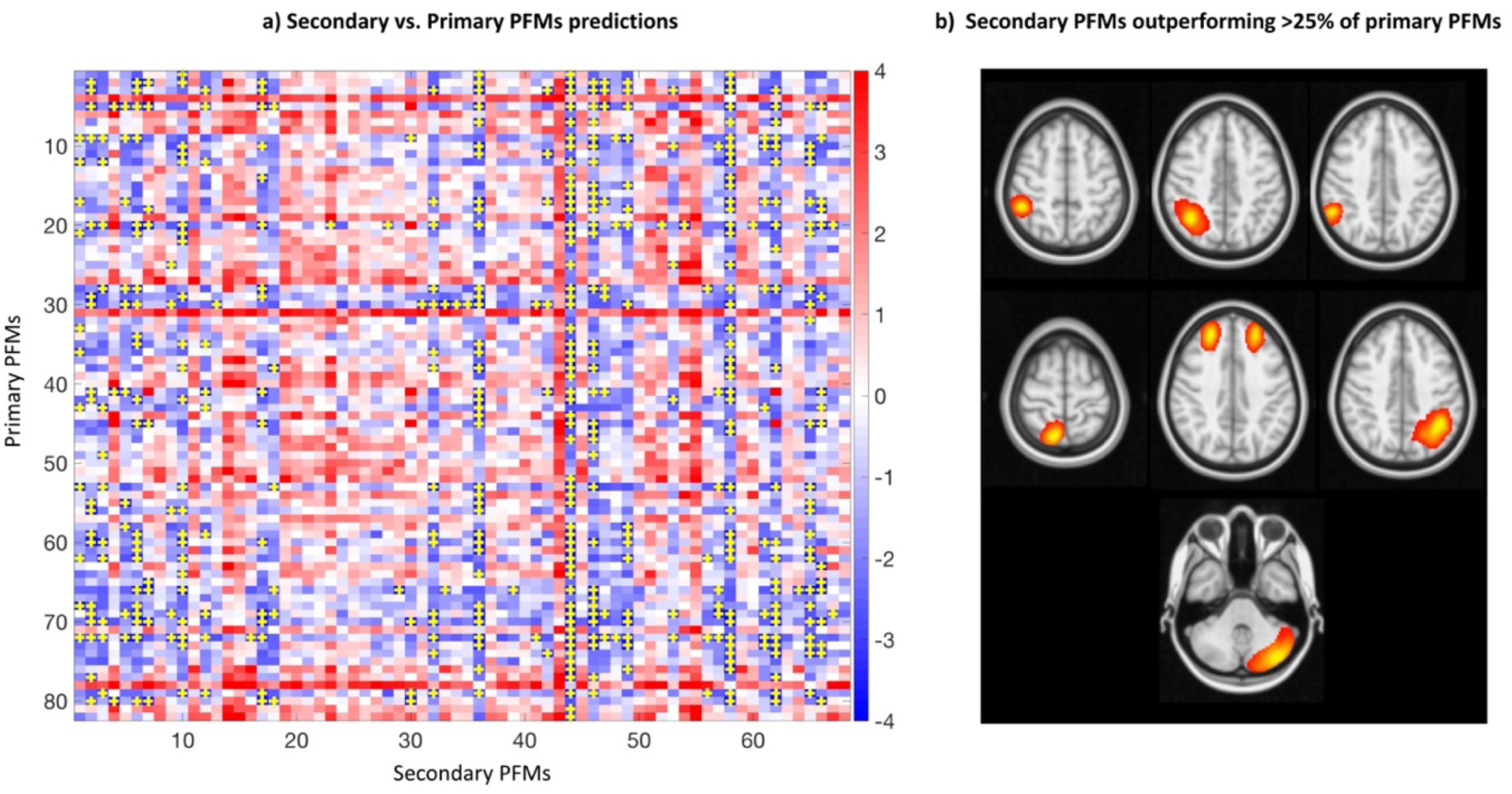
Comparing secondary PFMs to primary PFMs for prediction of cognitive tests. As outlined in 4.8, we categorised PFMs from 150-mode decomposition into two groups: 82 primary PFMs and 68 secondary PFMs. The latter are fine-grained modes that are found as we move to higher dimensions. a) t-values from weighted paired t-test comparison of multi-element prediction accuracies of every secondary PFM to all the primary PFMs. Blue shows primary<secondary, red vice versa. Locations marked with + denote secondary PFMs that are significantly better than primary PFMs, after FDR correction for multiple comparisons (across 68×82 tests). b) based on results in (a), seven secondary RSNs were found to outperform >25% of the primary RSNs for prediction of cognitive tests.

## Footnotes

1 We use functional mode (FM) as an umbrella term to describe both large-scale and parcel-like functional entities of the brain function in rest and task.

